# The structure of yeast Npl3 bound to RNA reveals a cooperative sequence-specific recognition and an RNA chaperone role in splicing

**DOI:** 10.1101/2022.05.24.493251

**Authors:** Ahmed Moursy, Antoine Cléry, Stefan Gerhardy, Katharina M. Betz, Sanjana Rao, Sébastien Campagne, Irene Beusch, Malgorzata M. Duszczyk, Mark D. Robinson, Vikram Govind Panse, Frédéric H.-T. Allain

## Abstract

The conserved SR-like protein Npl3 promotes splicing of diverse pre-mRNAs. However, the RNA sequence(s) recognized by the RNA Recognition Motifs (RRM1 & RRM2) of Npl3 during the splicing reaction remain elusive. Here, we developed a split-iCRAC approach in yeast to uncover the consensus sequence bound to each RRM. High-resolution NMR structures show that RRM2 recognizes a 5’-GNGG-3’ motif leading to an unusual mille-feuille topology. These structures also reveal how RRM1 preferentially interacts with a CC-dinucleotide upstream of this motif, and how the inter-RRM linker and the region C-terminal to RRM2 contributes to cooperative RNA-binding. Structure-guided functional studies show that Npl3 genetically interacts with U2 snRNP specific factors and we provide evidence that Npl3 melts U2 snRNA stem-loop I, a prerequisite for U2/U6 duplex formation within the catalytic center of the B^act^ spliceosomal complex. Thus, our findings provide insights into an unanticipated RNA chaperoning role for Npl3 during spliceosome active site formation.

## INTRODUCTION

Serine and Arginine-rich (SR) proteins belong to a family of proteins best known for their function in pre-mRNA splicing regulation, as well as multiple steps of gene expression from transcription to translation (Howard & Sanford, 2015, Jeong, 2017). SR proteins typically contain one or two N-terminal RNA-recognition motifs (RRM) followed by a C-terminal arginine-serine-rich (RS) domain (Manley & Krainer, 2010). In humans, 12 proteins belong to the SR protein family named SRSF1 to SRSF12 (Manley & Krainer, 2010). They are found in all metazoans in which they play a role in constitutive and alternative splicing regulation of most genes (Boucher, Ouzounis et al., 2001). In contrast, only around 4% of the genes of the budding yeast *Saccharomyces cerevisiae* contain introns and a few among them are alternatively spliced (Davis, Grate et al., 2000, Schreiber, Csaba et al., 2015). Consequently, only three SR-like proteins were identified in budding yeast (Gbp2, Hrb1, and Npl3) (Gilbert, Siebel et al., 2001, Hacker & Krebber, 2004). Of the three proteins, only Npl3 can promote splicing of intron-containing genes (Kress, Krogan et al., 2008). Npl3 was proposed to facilitate the splicing reaction by promoting the co-transcriptional recruitment of the spliceosome on chromatin through interactions via U1 and possibly U2 snRNPs (Kress et al., 2008) and with the Rad6 complex that adds a mono-ubiquitin to the histone H2B (Moehle, Ryan et al., 2012). Recently, Npl3 was also shown to be involved in the late steps of yeast spliceosome assembly by stimulating Prp28 helicase activity when Npl3 is phosphorylated. It was proposed that Npl3 may be the functional counterpart of the metazoan Prp28 N-terminal region that is absent in the yeast counterpart (Yeh, Chang et al., 2021). Finally, Npl3 was also shown to be required for the proper execution of the meiotic cell cycle by promoting splicing of introns containing non-consensus splice sites (Sandhu, Sinha et al., 2021). Npl3 has been implicated to function in multiple processes of gene expression including mRNA transcription elongation and termination (Bucheli & Buratowski, 2005, Bucheli, He et al., 2007, Dermody, Dreyfuss et al., 2008, Holmes, Tuck et al., 2015, Wong, Tang et al., 2010), mRNA export under stress (Gilbert et al., 2001, Hacker & Krebber, 2004, Shen, Stage-Zimmermann et al., 2000, Zander, Hackmann et al., 2016) and translation (Baierlein, Hackmann et al., 2013, Estrella, Wilkinson et al., 2009). Npl3 was also reported to maintain genome chromatin stability by preventing R-loops formation (Santos-Pereira, Herrero et al., 2013), contributing to telomere maintenance (Lee-Soety, Jones et al., 2012) and promoting double-strand DNA break repair (Colombo, Trovesi et al., 2017). Npl3 shares many functions with other metazoan SR proteins, and therefore serves as an ideal model to understand the evolution of SR proteins.

Npl3 is composed of two consecutive RRMs separated by a flexible eight amino acids linker that are followed by a C-terminal RS domain containing an Arg-Gly-Gly (RGG) repeat (Bossie, DeHoratius et al., 1992, Russell & Tollervey, 1992). The first canonical RRM is followed by a second pseudo-RRM (Birney, Kumar et al., 1993, Clery, Blatter et al., 2008). Although Npl3 binding to RNA is important for its functions, its mode of RNA-recognition remains elusive. The structures of the two RRMs of Npl3 were previously determined in their free form and showed that both RRMs do not interact (Deka, Bucheli et al., 2008, Skrisovska & Allain, 2008). While Npl3 was shown to bind preferentially to UG-rich RNA sequences using primarily its pseudo-RRM (RRM2) (Deka et al., 2008), the precise RNA sequence motifs recognized by the two RRMs remain unknown.

Here, we developed split-iCRAC, a combination of CRAC (Holmes et al., 2015) and iCLIP technologies (Huppertz, Attig et al., 2014), to identify the RNA sequences bound by yeast Npl3 *in vivo*. Using NMR spectroscopy, we determined the structures of both RRMs bound to a representative RNA consensus sequence obtained with the split-iCRAC approach. Our analyses revealed that RRM1 binds preferentially upstream of the RRM2 binding site. Both domains recognize a distinct RNA motif: 5’-NCCN-3’ and 5’-GNGGN-3’, respectively (N is for A, C, G or U) with the interdomain linker contributing to the RNA binding. Structure-guided studies revealed that mutations within RRM1, but not RRM2, negatively impact Npl3 function. However, Npl3 RRM2 had a specific effect on the splicing reaction, that is mediated through an unanticipated interaction of the protein with the U2 snRNA. We showed that this interaction destabilizes the U2 snRNA stem-loop I, thus providing mechanistic insights into an RNA chaperoning role for Npl3 during the formation of the spliceosome active site.

## RESULTS

### Identification of a consensus RNA motif recognized by Npl3 using iCRAC

Npl3 preferentially binds to U/G rich sequences *in vitro* (Deka et al., 2008) and *in vivo* (Holmes et al., 2015). However, the precise RNA motif(s) recognized by the two RRMs has remained elusive. To uncover the recognition motif of Npl3 *in vivo*, we performed a Crosslinking and Analysis of cDNA (CRAC) experiment with two key modifications (Holmes et al., 2015). First, we introduced an HRV-3C protease cleavage site directly after the sequence of RRM2 to distinguish RNAs that interact with the RRMs from those that are bound to the RGG/RS domain (Fig. 1A). A similar strategy was used to analyse exosome targets (Schneider, Kudla et al., 2012). Second, we used the individual-nucleotide resolution Cross-Linking and Immunoprecipitation (iCLIP) strategy to prepare the cDNA library and obtain single nucleotide resolution of the crosslinking sites (Huppertz et al., 2014). We termed this approach split-iCRAC. After HRV-3C cleavage, most of the RNA bound by Npl3 was detected with the RRMs (Fig. 1B), and a smaller fraction of RNA was detected bound to the RGG/RS domain. iCRAC libraries were then prepared with RNA isolated from the full-length protein (FL) and the RRM1/2 or RGG/RS domains. After mapping the sequencing reads to the *S. cerevisiae* genome, unique cDNA sequences from all replicates per sample were merged and used for cluster definition. *De novo* motif search was done using the HOMER software (Heinz, Benner et al., 2010) with sequences containing ±5 nucleotides around the identified clusters. Among all identified consensus motifs (Fig. S1A), only three were present in both FL and RRM1/2 and absent in the negative control (same experiment without the expression of tagged protein in cells) (Fig. S1B) leading to the determination of three consensus sequences, two for the FL protein and one for RRM1/2 (Fig. 1C). The three motifs have very similar sequences suggesting that the RGG/RS domain does not contribute to the specificity of RNA recognition. This finding is further supported by the absence of a specific motif enriched with the RGG/RS domain alone (Fig. S1). Moreover, the identified RNA bound by the RGG/RS domain are U-rich, which most likely reflects the higher efficiency of UV crosslinking to uracil over other nucleotides (Shetlar, Carbone et al., 1984).

**Figure 1:**
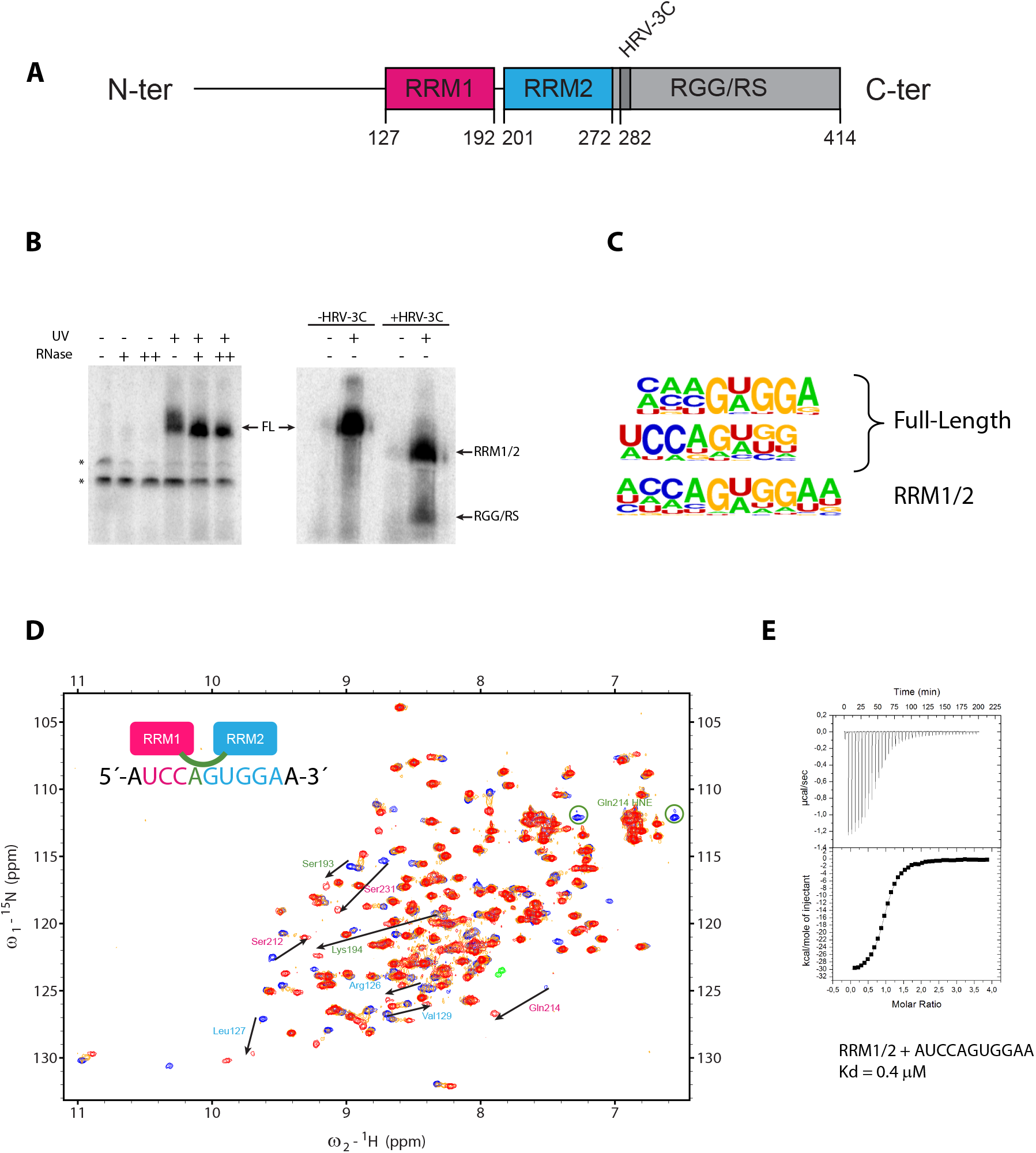
The split-iCRAC reveals the RNA motifs bound by Npl3 in vivo. **(A)** Schematic representation of Npl3 domain composition. The two RRMs are followed by an RS domain that is rich in RGG repeats. Amino acids at the border of each domain are numbered. The HRV-3C cleavage site inserted between the RRMs and the RGG/RS domain is shown. **(B)** Autoradiography of ^32^P labeled RNA after migration of the crosslinked complex on an SDS-PAGE gel. A negative control without exposure to UV shows no RNA band at the size of Npl3. Upon crosslinking with UV, the band becomes sensitive to RNase treatment. The stars represent unspecific bands in the initial experiments that were not observed anymore with the final optimized protocol. Treatment of the samples with HRV-3C protease separates the RNA bound to both RRMs from the RGG/RS domain. **(C)** Enriched motifs identified by split-iCRAC with Npl3 full length (top 2) and RRM1/2 using HOMER de novo motif finding on split-iCRAC derived clusters ±5 nucleotides. **(D)** Overlay of ^1^H-^15^N HSQC spectra recorded during the NMR titration of ^15^N labeled Npl3 RRM1/2 with increasing amount of unlabeled 5’-AUCCAGUGGAA-3’ RNA containing the bipartite motif identified by split-iCRAC (free form of the protein in blue, protein:RNA ratios of 1:0.3 and 1:1 in orange and red, respectively). Key residues of RRM1, RRM2 and the inter-domain linker for which a shift is observed upon RNA binding are indicated by an arrow labeled with pink, cyan and green colors, respectively. **(E)** ITC measurement performed with Npl3 RRM1/2 and the AUCCAGUGGAA RNA.

### Mode of interaction of Npl3 RRM1/2 with RNA

The split-iCRAC approach identified 5’-WCCAGWGGA-3’ (where W is U or A) as the consensus sequence interacting with RRM1/2 and the full-length Npl3 (Fig. 1C). To validate these findings, we monitored the binding of a recombinant Npl3 containing the two RRMs connected by their natural linker (RRM1/2, amino acids 114 to 282) to 5’-AUCCAGUGGAA-3’ RNA using NMR spectroscopy. Upon titration of the RRM1/2 construct by the RNA, several NMR chemical shift perturbations were observed for both RRM1 and RRM2 amide protons in the fast to intermediate exchange regimes (Fig. 1D). Saturation was reached at a 1:1 RNA:protein ratio indicating that one molecule of RNA was bound by both RRMs. The average correlation time measured for the complex was ∼13 ns, which corresponds to a size of about 22 kDa (Gossert & Jahnke, 2016, Rossi, Swapna et al., 2010). These data are consistent with molecular weights of 19 and 3 kDa for RRM1/2 and the RNA, respectively, and indicate that the two RRMs tumble with RNA as a single unit (Fig S2). A dissociation constant (K_d_) of 0.4 μM was determined with ITC for this complex (Fig. 1E). The identified consensus sequence contains a GG dinucleotide (Fig. 1C), which has been previously reported to be the common binding site of most known pseudo-RRMs, including Npl3 RRM2 (Clery, Sinha et al., 2013) since residues involved in the recognition of this dinucleotide are conserved in pseudo-RRMs (Fig 2A). Moreover, chemical shift perturbations observed with the isolated RRM2 of Npl3 and a sequence corresponding to the 3’ end of the RNA (5’-AGUGGAC-3’) (Fig. 2B) were very similar to those observed in the context of RRM1/2 bound to the longer RNA (Fig. S3). A K_d_ of 2 μM was determined with ITC for this complex (Fig. 2C). Taken together, this indicates that Npl3 RRM2 interacts with the 3’ extremity of the RNA, using a mode of RNA recognition common to all pseudo-RRMs (Clery et al., 2013).

**Figure 2:**
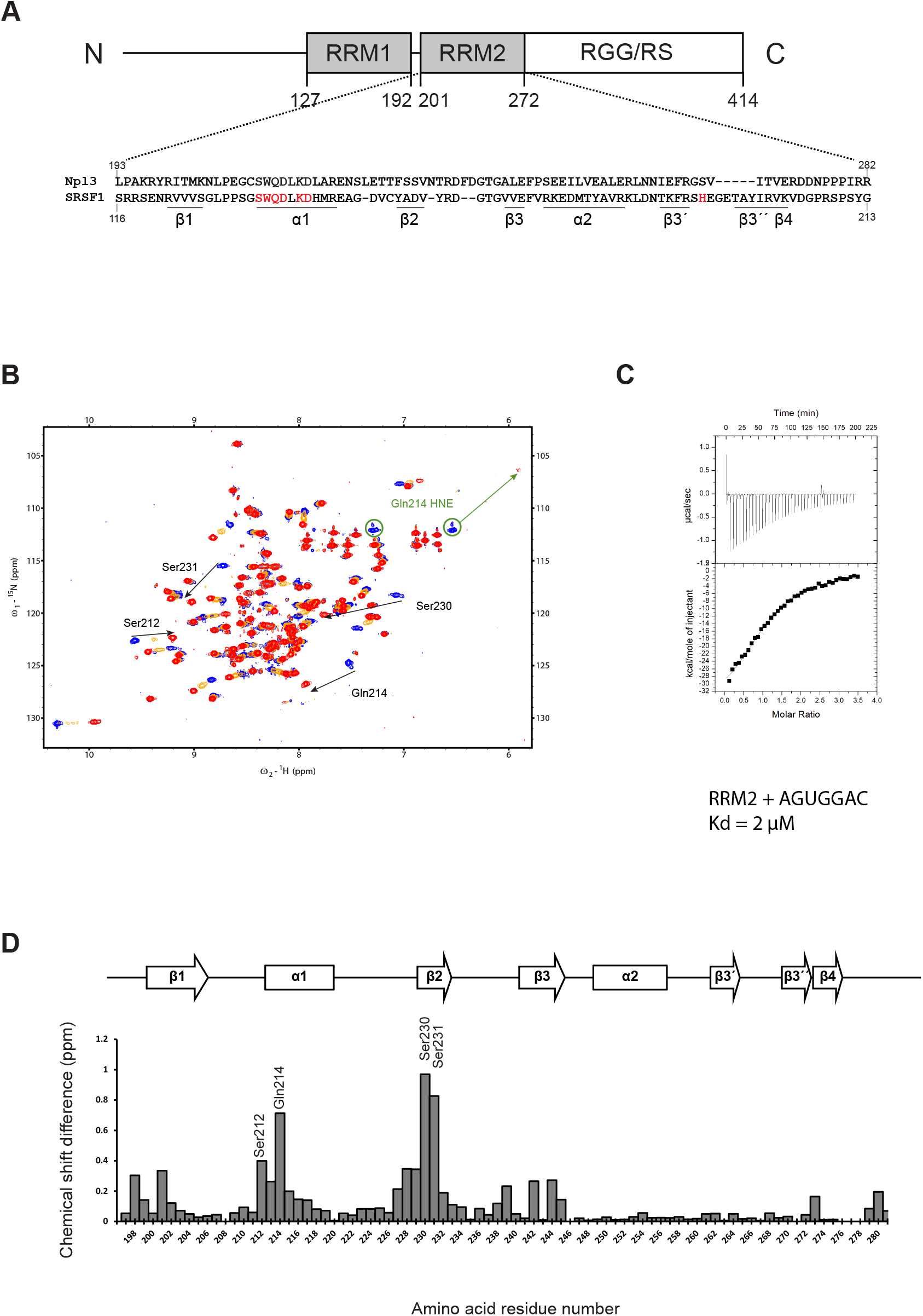
The specific interaction of Npl3 RRM2 with RNA. **(A)** Schematic representation of Npl3 domain composition. The sequence of Npl3 RRM2 is shown and aligned with the one of the RRM2 of human SRSF1. Residues that are important for SRSF1 RRM2 binding to RNA are colored in red. **(B)** Overlay of ^1^H-^15^N HSQC spectra recorded during the NMR titration of ^15^N labeled Npl3 RRM2 with increasing amount of unlabeled 5’-AGUGGAC-3’ RNA. The titration was performed at 40°C in the RRM2 NMR buffer. The peaks corresponding to the free and RNA-bound protein states (RNA:protein ratios of 0.3:1 and 1:1) are colored in blue, orange, and red, respectively. Black arrows indicate the most prominent chemical shift perturbations observed upon RNA binding. **(C)** ITC measurement performed with Npl3 RRM2 and the AGUGGAC RNA. **(D)** Representation of the combined chemical shift perturbations of Npl3 RRM2 amide residues upon binding to the 5’-AGUGGAC-3’ RNA at a ratio of 1:1 as a function of residue numbers. The corresponding secondary structure elements are represented at the top of the graph. The highest chemical shift perturbations annotated in **(B)** are indicated.

### Structure of Npl3 RRM1 bound to RNA

To elucidate the RNA-binding specificity of Npl3 RRM1 independently from RRM2, we performed NMR titrations of the isolated RRM1 with several 6mer ssDNA containing stretches of A, C, G or T as well as a 6mer polyU RNA (Fig. S4). Chemical shift perturbations were only detected with the polyC sequence indicating a strong preference of RRM1 for this nucleobase. We then used a modified version of the scaffold independent analysis (Beuth, Garcia-Mayoral et al., 2007) with ssDNA containing CX or XC motif (X is for A, C, G or T) flanked by degenerated sequences (Fig. 3A). We could then identify by NMR spectroscopy the motifs bound by Npl3 RRM1 with the highest affinity using the mean chemical shift perturbations observed upon ssDNA binding. As shown in Figure 3A, a clear preference for a CC dinucleotide over the other sequences was observed. Interestingly, the sequences selected with the split-iCRAC experiment were enriched in cytosines upstream of the motif bound by RRM2 (Fig. 1C), strongly suggesting that it corresponds to the RRM1 binding site. This result was rather unexpected, as Npl3 was never reported to bind preferentially to cytosines. Therefore, we investigated the interaction of the domain with the split-iCRAC derived RNA sequence 5’-AUCCAA-3’. A K_d_ value of about 15 μM was determined with ITC for this interaction (Fig. 3B). TOCSY experiments revealed that both cytosines are bound by different protein pockets (Fig. 3C) since two different chemical shifts were observed in their bound forms. Chemical shift perturbations of RRM1 bound to this short RNA (Fig. 3D, E) were very similar to those observed with RRM1/2 bound to the larger split-iCRAC RNA sequence (Fig. S3), indicating the functional relevance of using this small complex to characterize the mode of RNA recognition of Npl3 RRM1.

**Figure 3:**
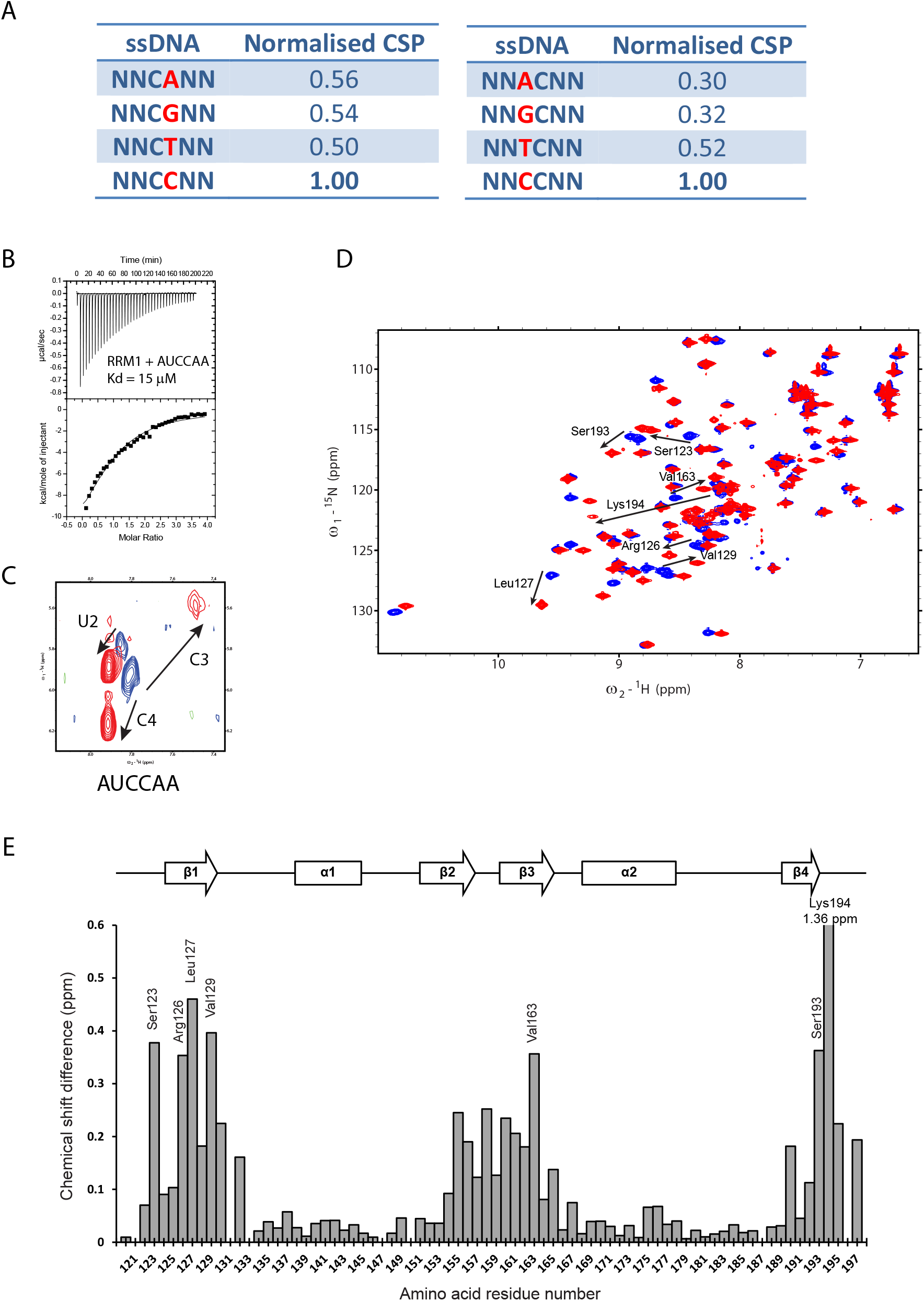
The specific interaction of Npl3 RRM1 with RNA. **(A)** Modified Scaffold independent analysis performed by titrating Npl3 RRM1 with 6mer ssDNAs. N is for any nucleotide (A, T, C or G). The normalized CSP represents the sum of combined chemical shift perturbations of non-overlapping peaks upon binding of the ssDNA to the RRM1 at a 1:1 ratio. The value was then normalized to the one obtained with the 5’-NNCCNN-3’ ssDNA. **(B)** ITC measurement performed with Npl3 RRM1 and the AUCCAA RNA. **(C)** Overlay of TOCSY spectra recorded with unlabeled RNA in the absence (in blue) and in the presence of Npl3 RRM1 at a 1:1 ratio (in red). Arrows represent the movement of the H5-H6 cross peaks for U2, C3, and C4 in different directions upon protein binding. **(D)** Overlay of ^1^H-^15^N HSQC spectra recorded during the NMR titration of ^15^N labeled Npl3 RRM1 with increasing amount of unlabeled 5’-AUCCAA-3’ RNA. The titration was performed at 40°C in the RRM1 NMR buffer. The peaks corresponding to the free and RNA-bound protein states (RNA:protein ratio of 1:1) are colored in blue and red, respectively. Black arrows indicate the most prominent chemical shift perturbations observed upon RNA binding. **(E)** Representation of the combined chemical shift perturbations of Npl3 RRM1 amide residues upon binding to the 5’-AUCCAA-3’ RNA at a ratio of 1:1 as a function of residue number. The corresponding secondary structure elements are represented at the top of the graph. The highest chemical shift perturbations annotated in (**D**) are indicated. The largest shift is observed for Lys194 with a value of 1.36 ppm.

We then determined the solution structure of RRM1 bound to 5’-AUCCAA-3’ using 2475 NOE-derived distance restraints including 135 intermolecular ones. We obtained a precise structure with an RMSD of 0.41 Å (Fig. 4A, Table S1). The RNA is lying on the surface of the RRM β-sheet with all nucleotides adopting an “*anti*” conformation and a C2’ endo sugar pucker conformation (Fig. 4B). Intermolecular contacts were observed between the U_2_, C_3_, C_4_, A_5_ and residues from the β-sheet and C-terminal extremity. U_2_ and A_5_ are not sequence-specifically recognized by the domain but provide binding affinity via their stacking on Arg130 side chain and C_4_ base, respectively. The C_3_ and C_4_ are sequence-specifically recognized by Npl3 RRM1. The C_3_ base stacks on Phe128 aromatic ring located on the β1-strand and forms two H-bonds between the amino and carbonyl groups of the base and the main chains of Tyr192 and Lys194, respectively (Fig. 4C). This latter H-bond is well supported by the fact that the amide of Lys194 experiences the largest chemical shift perturbation (1.36 ppm) upon binding to RNA (Fig. 3E). In addition, Lys194 from the C-terminus lies on top of the C_3_ base and its Lys194 amino group forms an H-bond with the 2’OH of C_3_. The C_4_ base stacks on Phe162 and is recognized by two H-bonds involving its O2 and N3 atoms and the side chain of Arg126. Finally, the aromatic ring of Phe160 contacts the riboses of both cytosines contributing to the binding affinity (Fig. 4C). Based on these structural data, we conclude that Npl3 RRM1 recognizes a 5’-NCCN-3’ motif.

**Figure 4:**
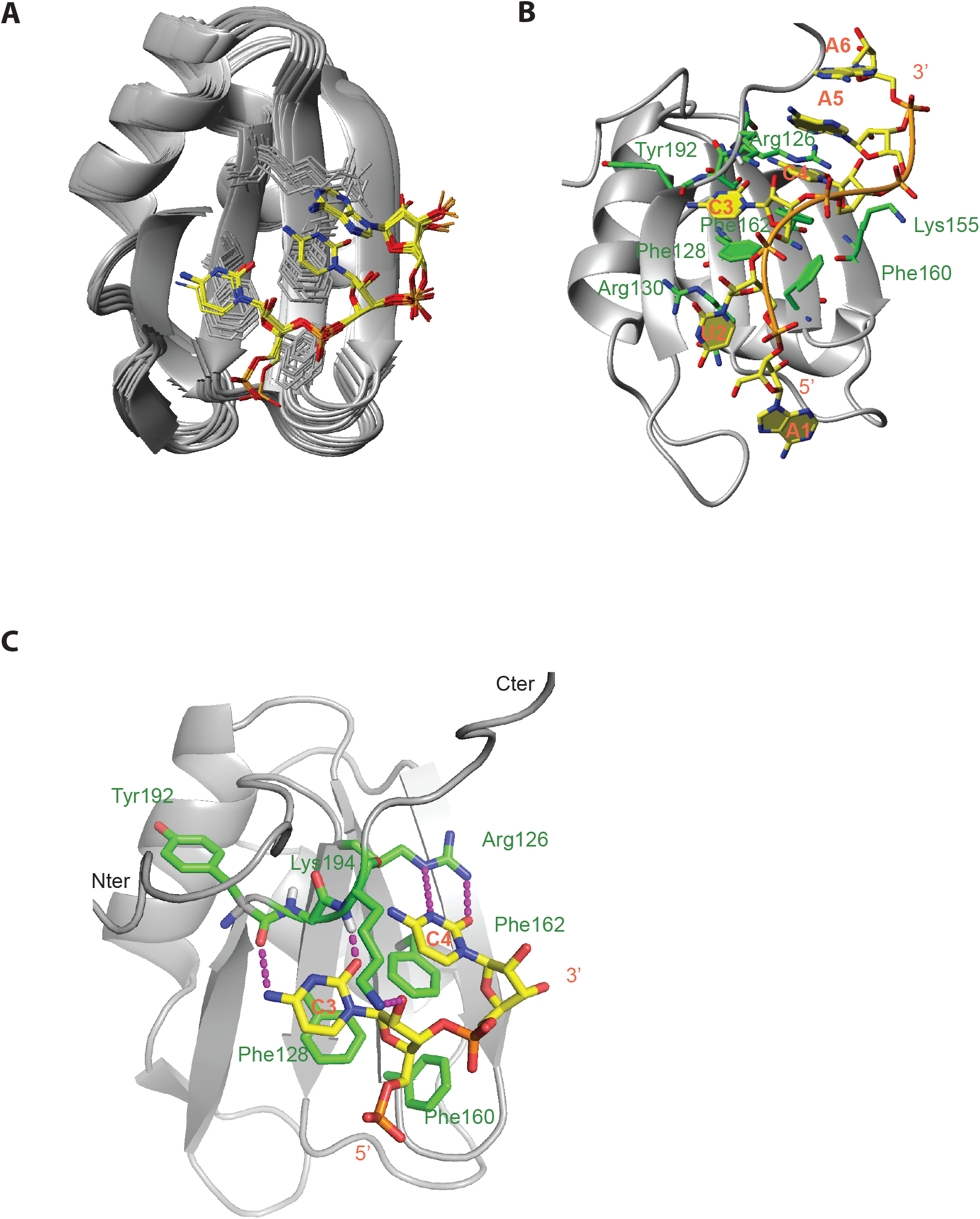
Overview of the solution structure of Npl3 RRM1 bound to the 5’-AUCCAA-3’ RNA. **(A)** Overlay of the 10 lowest-energy structures superimposed on the backbone of the structured parts of the protein and heavy atoms of RNA. The protein backbone is shown in grey and heavy atoms are shown in orange (P atoms), yellow (C atoms of RNA), red (O atoms) and blue (N atoms). The RRM (residues 120 to 198) and the ordered region of RNA (C3, C4, A5) are shown. **(B)** The solution structure of the complex is shown in ribbon (protein backbone) and stick (RNA) representation. Protein side-chains or backbone involved in RNA interactions are shown as sticks. C atoms of the protein are in green. **(C)** Details of the RNA recognition by Npl3 RRM1. H-bonds are in magenta.

### Structure of Npl3 RRM1/2 bound to RNA

The affinity of RRM1/2 for RNA (K_d_ = 0.4 μM) was significantly higher than each isolated RRM (K_d_ values of 15 and 2 μM, respectively) (Figs. 1, 2 and 3) suggesting a cooperative mode of interaction of the two domains with RNA and/or additional contacts mediated by the inter-domain linker. The binding of RRM1 to the 5’-NCCN-3’ motif indicated an interaction of RRM1 with RNA upstream of the RRM2 binding site on the split-iCRAC defined sequence. To investigate whether the orientation of the two RRMs was important for the binding efficiency of Npl3 to RNA, we performed an NMR titration of RRM1/2 with the 5’-AUGGAGUCCAA-3’ RNA containing inverted binding motifs (RRM2 binding site at the 5’-end and RRM1 binding motif at the 3’-end). As illustrated in Figure S5A, smaller CSPs were consistently observed at saturation (1:1 protein:RNA ratio) with this RNA compared to the 5’-AUCCAGUGGAA-3’ RNA, showing that the binding of Npl3 RRM1/2 to this RNA is apparently weaker than with the split-iCRAC derived sequence. Accordingly, the affinity measured for this complex by ITC indicates a K_d_ of 1.2 μM (Fig. S5B) which is 3 times weaker than with the 5’-AUCCAGUGGAA-3’ RNA (Fig. 1E). Although both RRMs are bound to this RNA with RRM2 binding upstream and RRM1 binding downstream, these data suggest that for optimal RNA binding, RRM1 and RRM2 should bind their respective sequence upstream and downstream, respectively.

Next, we investigated the mode of RNA recognition of both Npl3 RRMs. Although many intermolecular NOEs between Npl3 RRM1/2 and 5’-AUCCAGUGGAA-3’ could be observed, the quality of the NMR data was always better in the complexes with isolated RRMs. However, the similarity of the chemical shift perturbations observed upon RNA binding for the isolated RRMs and the RRM1/2 complex (Fig. S3) indicated that the mode of interaction of Npl3 RRMs was identical in both cases. In addition, most intermolecular NOEs found in the complexes with the isolated RRMs were also observed with RRM1/2-RNA complex confirming the same mode of interaction of the domains with RNA in both contexts. Therefore, to calculate the structure of the RRM1/2 complex, we used the same intermolecular NOEs as in each single RRM complex although some of them were too broad to be observed with the larger Npl3 RRM1/2 complex. We used this strategy only when the intermolecular contacts were confirmed by similar chemical shift perturbations. Additionally, due to an unfavorable exchange condition, we could not detect any intermolecular NOEs between G9 imino and RRM2 in any complexes. However, in our preliminary structures, the position of the G9 is very similar to the equivalent guanine in SRSF1 RRM2-GGA complex but less precisely defined due to the missing intermolecular constraints (Clery et al., 2013). In order to more precisely position this base, we used in our structure calculations the same restraints for G9 H1 and residues of Npl3 RRM2 as for the structure determination of SRSF1 RRM2 bound to RNA (Clery et al., 2013). Those restraints did not induce any distance violations indicating that they were in perfect agreement with all other experimentally derived restraints.

In total, to calculate the structure of RRM1/2 bound to the 5’-AUCCAGUGGAA-3’ RNA, we used 3788 distance restraints including 189 intermolecular ones and 62 Residual Dipolar Coupling (RDC) derived restraints (Table S2). We could reach a high precision with a heavy atom RMSD of 1.23 Å (Fig. 5A). The two RRMs are precisely positioned relative to each other due to an interaction of Npl3 RRM2 with G_6_ that was not present in the complex formed with the pseudo-RRM in isolation or in SRSF1 RRM2 complex. This nucleotide identity of G_6_ is strictly conserved or largely dominant in the split-iCRAC consensus sequences (Fig. 1C). G_6_ adopts a *syn* conformation and stacks on the Phe229 aromatic ring located on RRM2 β2-strand. G_6_ is also contacted by the C-terminal region of RRM2, with the side-chain of Ile 279 contacting its base and sugar (8 intermolecular NOEs between these two residues position the C-terminal end near the RNA and RRM1). These contacts also explain the chemical shift changes seen for Ile 279 and Arg280 upon RNA binding (Fig. S3). In fact, in some of the structural conformers, Arg280 and Arg281 are positioned sufficiently close of RRM2 to interact via a salt bridge with the side chains of Glu 153 and Glu 138. These additional protein-protein contacts help rationalizing why the two RRMs adopt a fixed orientation upon RNA binding (Fig. 5C).

**Figure 5:**
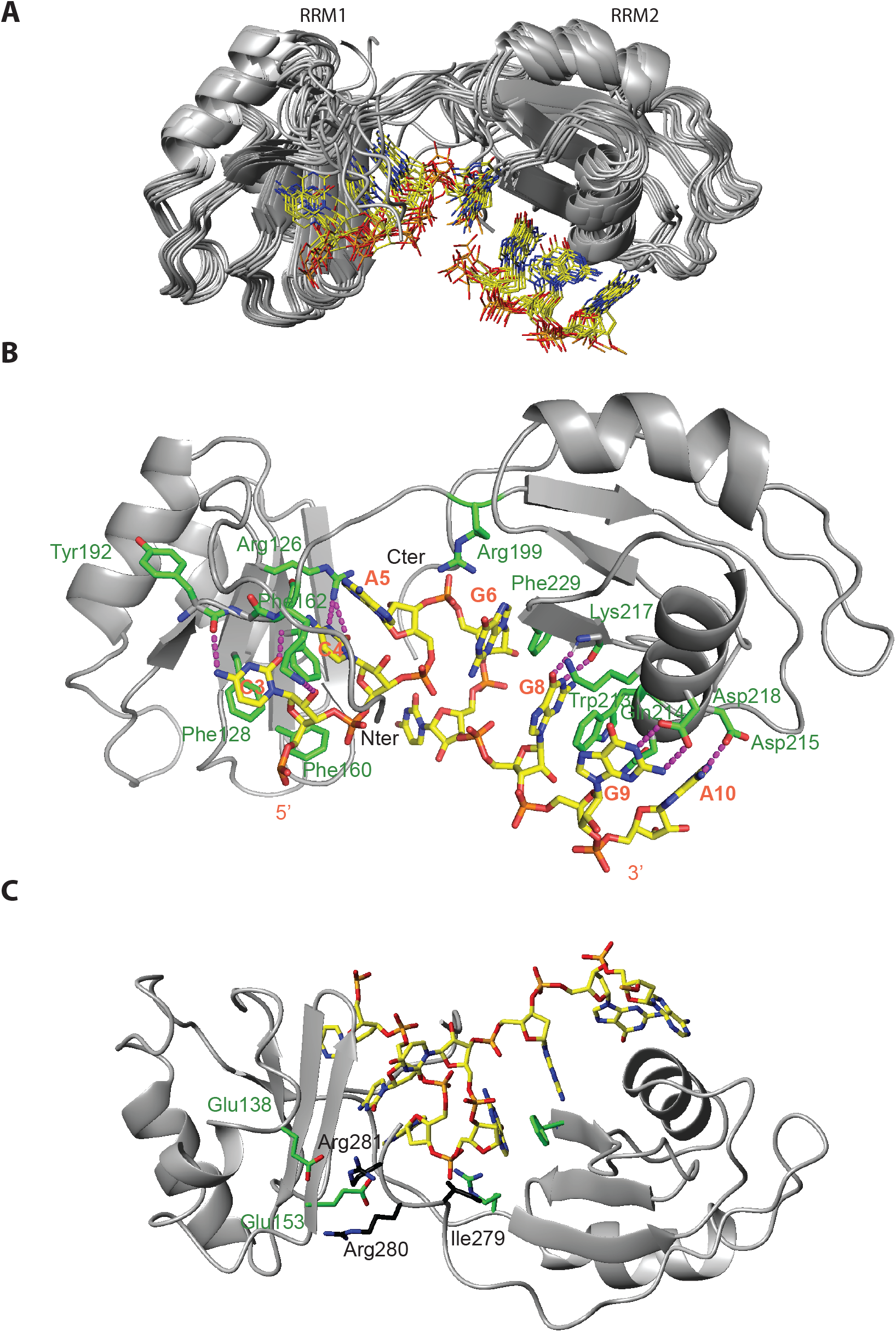
Overview of the structure of Npl3 RRM1/2 bound to the 5’-AUCCAGUGGAA-3’ RNA. **(A)** Overlay of the 10 lowest-energy structures superimposed on the backbone of the structured parts of the protein and heavy atoms of RNA. The protein backbone is shown in grey and heavy atoms are shown in orange (P atoms), yellow (C atoms of RNA), red (O atoms) and blue (N atoms). The RRMs (residues 12 to 198) and the ordered region of RNA (C3 to G6 and G8 to A10) are shown. **(B)** The solution structure of the complex is shown in ribbon (protein backbone) and stick (RNA) representation. Protein side-chains or backbone involved in RNA interactions are shown as sticks. C atoms of the protein are in green. H-bonds are in magenta. **(C)** The solution structure of the complex is shown from the back with the residues of the C-terminal part of the protein (Ile279, Arg280 and Arg281) involved in interactions with G_6_ and the Glu138 and Glu153 residues of RRM1.

Although no inter-NOE could be observed between the Arg199 and the base, the structure suggests that this side chain may interact with G_6_ explaining the specific recognition of a guanine at this position (Fig. 5C). In good agreement, the Arg199 NH_ε_ disappears upon complex formation. Furthermore, the chemical shift of G_6_ H8 was shifted in the RRM1/2 complex compared to the complex with the isolated RRM2 indicating that this proton is in a different environment when the two RRMs are bound. Overall, the GNGGA motif is tightly bound by RRM2 via a series of six consecutive stacking interactions (G_6_/ Phe229/G_8_/Trp213/Gln214/A_10_) adopting a “mille-feuille” topology, which certainly contributes to the higher RNA affinity of RRM2 compared to RRM1.

To investigate the importance of the protein-RNA contacts observed in these structures, we measured the binding affinity of several Npl3 RRM1/2 alanine mutants of key residues involved in RRM1 or RRM2 interaction with RNA. Mutations of residues that are important for the specific recognition of the CC dinucleotide binding by RRM1 (R126A and F128A) decreased the binding affinity strongly from a K_d_ of 0.4 μM to 13 and 12 μM, respectively (Fig. S6). The mutation of R130 that stacks underneath the U_2_ base had a milder effect on binding affinity (K_d_ of 1.1 μM). Mutations that affect the recognition of the GG dinucleotide by RRM2 (Q214A and K217A) also showed a moderate decrease in binding affinity with K_d_ values increasing to 1.5 and 7 μM, respectively (Fig. S6). As expected, the protein variant carrying both mutations showed weaker binding (Kd of 16 μM). Interestingly, the mutation of Phe229 that contacts G_6_ had also a clear effect on the RNA binding affinity (K_d_ of 3.8 μM). The importance of this interaction was further corroborated by the drop in binding affinity (K_d_ of 2.7 μM) observed between WT RRM1/2 and a G_6_A RNA variant (Fig S7). Additional mutations of the RNA were tested, including the conversion of C_3_ or C_4_ to uracil, which decreased the binding affinity to a K_d_ value of 1.3 and 1.2 μM, respectively. G_8_ or G_9_ mutation to adenine resulted in a slightly higher affinity drop (K_d_ of 2μM). Surprisingly, shortening the RNA by removing A_5_ did not affect RRM1/2 binding affinity. These data indicate a certain flexibility regarding the position of the two RRMs relative to each other, allowing them to adapt their binding mode to a larger repertoire of RNA sequences. In agreement with these *in vitro* data, these two possible binding registers are seen in the Npl3 first split-iCRAC consensus motifs selected with the full-length protein were A and C nucleotides are equally present before the conserved GNGG motif (Fig. 1C).

### Contribution of each RRM toward Npl3 function *in vivo*

Our *in vitro* investigation of Npl3 RRM1/2 interaction with RNA permitted to rationally design protein mutants that strongly decrease the binding of RNA with either RRM1 (F128A, F128A+F160A) or RRM2 (Q214A, K217A, Q214A+K217A) without affecting the folding of the protein (Fig. S8). These substitutions were used to investigate the impact of the RNA interaction of each RRM on Npl3 functionality *in vivo*. Yeast strains lacking the *npl3* gene (*npl3Δ*) were transformed with plasmids expressing these mutants under the control of the natural Npl3 promoter. We investigated whether they could rescue the slow growth of the *npl3Δ* strain (Kress et al., 2008). Interestingly, the complementation with Npl3 having single or double substitutions in RRM1 failed to completely rescue the slow growth defects (Fig. 6A) indicating the importance of RRM1 binding to RNA for Npl3 functionality. Conversely, single and double substitutions within RRM2 did rescue the Npl3 deletion phenotype (Fig. 6A) indicating a more critical contribution of the RRM1 RNA binding interface than RRM2 in Npl3 function. Note that the effect is not due to a difference in binding affinity since both double mutants have a similar K_d_ (Fig. S6).

**Figure 6:**
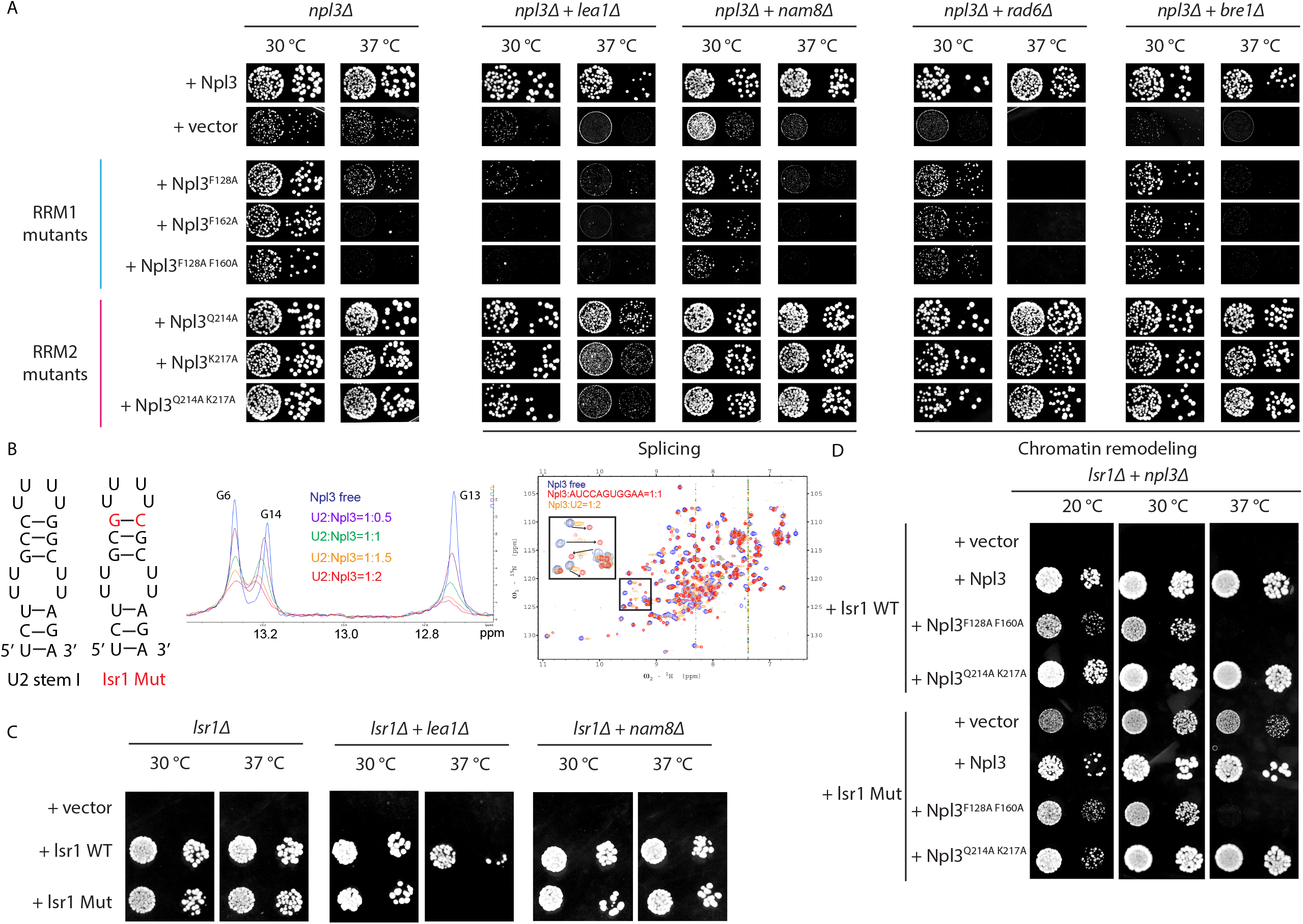
RRM1 and RRM2 are non-equivalently important for the functions of Npl3 in vivo. **(A)** Mutant growth analysis of npl3Δ strain complemented with vectors expressing different protein variants of Npl3 (marked on the left). Yeast cells were plated on SD-leu plates and incubated at 30 or 37 °C. Two steps of a yeast serial dilution are shown for each condition. Synthetic growth analysis of npl3Δ+lea1Δ, npl3Δ+nam8Δ, npl3Δ+rad6Δ and npl3Δ+bre1Δ double deletion strains complemented with Npl3 protein variants reducing the interaction of RRM1 (F128A, F162A, F128A+F160A) or RRM2 (Q214A, K217A, Q214A+K217A) with RNA is shown. Lea1 and Nam8 are involved in splicing, whereas Rad6 and Bre1 are chromatin-remodelling factors. **(B)** Sequence of yeast U2 snRNA stem I. The Isr1 mutation is shown in red. Overlay of 1D NMR spectra recorded at 303 K with U2 SL I free form and at U2 SL I:Npl3 RRM1/2 ratios of 1:0.5, 1:1, 1:1.5 and 1:2. Overlay of ^1^H-^15^N HSQC NMR spectra recorded with Npl3 RRM1/2 free form (blue), Npl3 RRM1/2:U2 SL I at a 1:2 ratio (orange) and :1 ratio (red). **(C)** Synthetic growth analysis of Isr1Δ, Isr1Δ+lea1Δ and Isr1Δ +nam8Δ deletion strains complemented with the WT or Mut versions of Isr1. **(D)** Synthetic growth analysis of the ΔIsr1+npl3Δ double deletion strain complemented with either the WT or Mut version of Isr1 and the same Npl3 protein double variants as tested in (**A**).

The slow growth phenotype of *npl3Δ* yeast mutant was previously reported to be exacerbated when combined with deletion of genes involved in spliceosome assembly (*lea1 and nam8*) (Kress et al., 2008) or chromatin remodelling (ex. *rad6* and *bre1*) (Moehle et al., 2012). We investigated the respective involvement of both Npl3 RRMs in these two functions by testing whether the RRM1 and RRM2 mutants of Npl3 would lead to the same genetic interaction. In agreement with the predominant importance of the RRM1 RNA binding interface, we observed genetic interactions of RRM1 mutated Npl3 with the four tested genes (Fig. 6A). This indicates that the binding of RRM1 to RNA was important for both the splicing and the interaction of Npl3 with chromatin remodelling factors. On the other hand, RRM2 mutants only showed a genetic interaction with *lea1* with a predominant effect seen at 37°C (Fig. 6A). Surprisingly, no effect was observed with *nam8* despite the involvement of both *lea1* and *nam8* in splicing regulation. Lea1 is part of the U2 snRNP, while Nam8 is a component of the U1 snRNP, indicating that RRM2 binding to RNA might be more linked to U2 snRNP function during splicing.

In good agreement with this hypothesis, Npl3 crosslinks were enriched at the 5’ end of U2 snRNA but not in U1 snRNA (Fig. S9). In addition, a sequence containing a CC followed by a GG dinucleotide (reminiscent of the motif recognized by both RRMs of Npl3), is found in this cross-linked region (Fig. S9A). Interestingly, those nucleotides are part of the U2 snRNA stem-loop I (SL I) in which the CC and GG are base-paired (Fig. 6B). However, in *S. cerevisiae*, this stem must be melted during the splicing reaction to allow the formation of a duplex between U2 and U6 snRNAs (Yan, Wan et al., 2016). Our structure clearly indicates that Npl3 RRMs interact sequence-specifically with single-stranded RNA at the CC (RRM1) and GG (RRM2) sequences (Fig. 4 and 5) suggesting that the binding of Npl3 to U2 snRNA SL I would induce the melting of this stem-loop. We then *in vitro* transcribed the 5’-GAG<u>CGAA**UCUCUUU**</u>**GCCUUUUG<u>GCUUAGA</u>**UC-3’ RNA containing the initiator codon GAG followed by the sequence forming the U2 stem-loop I (in bold) including the two parts involved in the duplex formation with the U6 snRNA (underlined sequences). Using NMR spectroscopy, we confirmed that this RNA adopted the expected secondary structure based on previous NMR assignments obtained with this stem-loop (Sashital, Venditti et al., 2007). In addition, the titration of this stem-loop with Npl3 RRM1/2 showed that the protein could interact with the U2 SL I sequence and destabilized the stem (Fig. 6B). Indeed, the intensity of the imino signals observed when the upper part of the stem is formed (G6, G13 and G14) decreased upon protein binding without any chemical shift perturbations indicating that the protein did not interact with the stem but rather unfolds it (Fig. 6B). The overlay of the 2D ^1^H-^15^N HSQC spectra recorded with Npl3 RRM1/2 in the free form and bound to the U2 SL I (Fig. 6B) validated this interaction. We therefore mutated the CC and GG binding sites to CG to keep the ability to form the stem I but prevent the potential binding of Npl3 to this RNA (Isr1 mut, Fig. 6B) and tested the effect in yeast. The mutation of U2 snRNA did not show any phenotype at any of the tested temperatures (Fig 6C). However, when combining the mutation with a deletion of the U2 factor Lea1, we could observe a slow growth phenotype at 30°C and complete lethality of the cells at 37°C (Fig 6C). This effect was not seen when combining the same mutation with a deletion of the U1 snRNP component Nam8 (Fig 6C). This indicated that the putative binding sequence of Npl3 identified in the stem-loop was important for the function of U2 snRNP. To confirm the link with Npl3, we tested a combination of mutations in Npl3 RRM1 or RRM2 with the Isr1 mutant. Interestingly, we observed a slow growth phenotype at lower temperature (20°C) confirming the link between Npl3 RRM2 and the U2 snRNA (Fig 6D). Overall, these data indicate that Npl3 favors the U2-U6 duplex formation required for the formation of the spliceosome active site by interacting with and destabilizing the stem-loop I of U2 snRNA.

## DISCUSSION

### Split-iCRAC reveals the RNA motif(s) recognized Npl3 *in vivo*

Despite the pivotal role of Npl3 in RNA metabolism, no RNA binding consensus sequence was identified for this protein yet. Only a preference for this protein in binding to UG-rich RNA sequences was previously reported (Deka et al., 2008, Holmes et al., 2015). Using a modified version of the CRAC method, we identified a clear consensus RNA sequence bound to yeast Npl3. Adding the individual nucleotide resolution of the iCLIP protocol allowed us to identify a consensus motif, which could not be determined using the conventional CRAC (Holmes et al., 2015). In general, the binding profile that we observed correlates well with the available CRAC data (Fig. S10). Interestingly, the consensus RNA sequence obtained with Npl3 (UCCAGUGGA) is different from the motifs identified by PAR-CLIP with the two other SR-like proteins Hrb1 (CuGCU) and Gbp2 (GGUG) (Baejen, Torkler et al., 2014) indicating that these proteins have distinct RNA targets in vivo. Hrb1 and Gbp2 contain three RRMs. Their most C-terminal RRM was shown to interact directly with proteins of the THO/TREX complex instead of binding to RNA (Martinez-Lumbreras et al., 2015). Their two N-terminal domains are involved in RNA binding but seem to recognize a shorter motif than Npl3. Like in Npl3, their second RRM is a pseudo-RRM. This domain was shown to be responsible for the recognition of the GGUG motif by Gbp2, which seems to use the same mode of recognition as described for the human SRSF1 pseudo-RRM (Clery et al., 2013) and Npl3 (this study) confirming the conservation of the mode of interaction of all pseudo-RRMs with two consecutive guanines (Clery et al., 2013). Nevertheless, these guanines were not present in the PAR-CLIP consensus sequence identified with Hrb1, which could indicate a predominant function of the canonical RRM1 in binding to the CuGCU motif.

Moreover, the “split” version of the iCRAC revealed that the two RRMs but not the RGG/RS domain were responsible for the specific interaction of Npl3 with RNA (Fig. 1B). Previously, split-CRAC was employed to study the binding of each RNA binding domain of Rrp44 to RNA by inserting a protease cleavage site in the interdomain linker (Schneider et al., 2012). The same strategy could not be used to separate the RNAs bound by RRM1 and RRM2 of Npl3 as the two domains are only eight amino acids apart, and some of those residues were shown to be involved in RNA binding (Fig. 5). Mutating those amino acids or elongating the linker to insert a protease recognition sequence might have influenced the RNA binding properties of the RRMs leading to false interpretations of the resulting data. Importantly, our split-iCRAC strategy also permitted the identification of RNA sequences that were directly bound by the RGG/RS domain. This domain generally binds RNA sequences near the binding sites of the RRMs with no apparent sequence preference either upstream or downstream (Fig. S10B). We could not detect any specific RNA motif bound by the RGG/RS domain apart from a general uracil enrichment, which most likely originates from a UV-crosslinking bias (Fig. S1). This result is in good agreement with recent reports showing that the RS domains bind non-specifically to RNA (Castello, Fischer et al., 2012, Jarvelin, Noerenberg et al., 2016). Another study reported that RS domains of SR proteins could bind directly to RNA sequences containing the splicing branch point (Shen & Green, 2004, Shen, Kan et al., 2004). However, the split-iCRAC data did not reveal any specific cross-links of the Npl3 RGG/RS domain around branch point sequences.

### Molecular basis of RNA recognition by Npl3 RRMs

Npl3 was previously shown to interact with UG-rich RNA sequences (Deka et al., 2008, Holmes et al., 2015) and no specific interaction for Npl3 RRM1 with RNA was reported so far. Here, we show that Npl3 RRM1 participates to the specific interaction of the protein with RNA by binding to CC dinucleotides (Fig. 4). Interestingly, the recognition of two consecutive cytosines was also observed in the structure of the human SR protein SRSF2 bound to RNA (Daubner, Clery et al., 2012). Despite the fact that the two RRMs are only 43% homologous (and 28% identical), the position of the two cytosines on the β-sheet surface and their recognition by the RRM are quite similar (Fig. S11A). However, unlike the SRSF2 RRM that recognizes CC, CG, GC and GG, Npl3 RRM favors strictly CC (Fig. 3A).

In addition, we found that RRM2 recognizes a 5’-GNGGA-3’ motif in the context of RRM1/2. The recognition of the GG dinucleotide is identical to what was reported for SRSF1 RRM2 (Clery et al., 2013). Although pseudo-RRMs share the recognition of a GG dinucleotide, the nucleotides bound on each side of this motif seem to be more specific to each protein. For instance, the adenine bound by SRSF1 downstream of the GG is positioned similarly in Npl3 RRM2. Nevertheless, the stacking interaction of A_10_ with the His193 observed in the structure of the SRSF1 complex is not possible with Npl3 as the β3-β4 hairpin is shorter and the corresponding aromatic residue is missing (Fig. 2A and 5). Another difference between the two complexes is the binding of Npl3 RRM2 to G_6_ upstream of the GG dinucleotide motif (Fig. 5). Side-chains from RRM2 (Phe 229), the interdomain linker (Arg 199) and the region C-terminal to RRM2 (Ile 279) contribute the specific recognition of this nucleotide.

RRM2 of Npl3 was previously reported to bind GU rich RNA sequences (Deka et al., 2008). In good agreement, our split-iCRAC motif showed that the sequence bound by RRM2 can contain uracils, as the consensus motif was GU_/A_GG (Fig 1C). In the structure of RRM1/2 bound to RNA, the uracil is bulged out (Fig 5) and does not give any intermolecular NOEs either in this complex or in the isolated domain bound to the 5’-AGUGGAC-3’ RNA. However, we observed that in the absence of G_2_ in the 5’-AUGGAC-3’ RNA, additional chemical shift perturbations were observed in the β2-β3 loop of RRM2. Those additional chemical shift perturbations were not observed with the AAGGUC RNA, which hints towards a possible specific recognition of the U_2_ 5’ to the GG dinucleotide. In this context, we could observe intermolecular NOEs between U_2_ and Val232 and Asn233 from the β2-β3 loop indicating that the uracil is indeed in contact with the protein. However, because of limited spectral quality, we could not precisely position the uracil to infer its specific recognition. This result suggests that RRM2 could either bind to GNGGA or UGGA motifs.

### Tandem RRMs bind an extended single-strand RNA via an unusual orientation

In addition to defining the exact RNA motifs recogniszed by RRM1 and RRM2, our structural studies revealed their relative orientation. Although no contact could be observed between the two RRMs of Npl3 in their free form (Deka et al., 2008, Skrisovska & Allain, 2008), their binding to a single RNA molecule was expected to rigidify the orientation of one relative to the other as reported previously with other tandem RRMs (Afroz, Cienikova et al., 2015). The preferential binding of RRM1 upstream of the RRM2 binding site is not common among tandem RRMs. In most structures, RRM2 binds RNA upstream of the RRM1 binding site (Afroz et al., 2015). The only case reported so far of two RRMs adopting an opposite orientation on RNA was with the tandem RRMs of TDP-43 (Fig. S11B) (Lukavsky, Daujotyte et al., 2013). To keep this unusual orientation, it was hypothesized that a long inter-domain linker (15 aa) is required to allow the two RRMs to lie side-by-side (β2-β4 type) and form an extended β-sheet RNA binding surface (Afroz et al., 2015). Recently, two structures (Dnd1 (Duszczyk, Wischnewski et al., 2021) and Npl3 in this work) revealed new ways for tandem RRMs to bind RNA cooperatively with RRM1 binding to the 5’end despite having a short interdomain linker (5 and 7 aa, respectively). In such cases, the canonical β-sheet surface of the RRM is not used but rather both sides of the domain. In Dnd1, the α2−β4 edge of the RRM is used while in Npl3, this is the α1−β2 edge (Fig. S11B). Having a short interdomain linker presents the advantage to reduce the entropy cost upon RNA binding. This illustrates once more the unusual diversity and rather unpredictable binding mode of RRM-RNA interactions (Afroz et al., 2015, Clery et al., 2008, Maris, Dominguez et al., 2005).

Another similarity between the structure of TDP43 and Npl3 is the presence of a guanine located in the centre of the bound RNA sequence, which is sequence-specifically recognized by both proteins and contributes to establish a fixed orientation of the two RRMs. However, the base adopts an *anti* conformation in the TDP-43 complex, whereas the base is *syn* when bound to Npl3. Surprisingly, despite the fixed orientation of RRMs on RNA, the RGG/RS domain of Npl3 seems to bind non-specifically upstream or downstream of their binding sites (Fig. S10B), suggesting a high flexibility of the domain. In addition, it raises an intriguing possibility that Npl3 recruits additional proteins on both sides of its binding site via this disordered region.

### Functional insights into the role of Npl3 during RNA splicing

Our genetic interactions implicate that the RRM1-RNA interactions are broadly important for the role of Npl3 during splicing and chromatin remodelling, whereas the RRM2 involvement seems to be rather limited to the splicing process (Fig 6A). The observed genetic interactions were more obvious when yeast cells were grown at 37 °C (Fig. 6A). In addition, the genetic interaction between RRM2 mutated *npl3* and *lea1* was only observed at 37 °C (Fig. 6A). One explanation is that at a higher temperature, the lower affinity of the RRM mutants for RNA is thermodynamically disfavored while it can still occur to some extent at lower temperatures.

Previous mutational analyses to study the function of Npl3 RRM2 *in vivo* used three mutations (L225S, G241N, and E244K) (Bucheli & Buratowski, 2005) mostly in combination (Colombo et al., 2017, Lee-Soety et al., 2012). However, it was previously reported that the L225S unfolds the RRM2 (Deka et al., 2008). The effect of the G241N and E244K on the folding of the RRM was never tested, but these two residues are far away from the RNA binding interface. Therefore, it is difficult to correlate a loss of function from those mutants with the RNA binding properties of RRM2. Similarly, *in vivo* mutational studies were previously done using the F160L mutation to investigate the function of RRM1 (Lee-Soety et al., 2012). However, the effect of this single mutation on RNA binding was never directly tested. Our structure shows that indeed Phe160 has hydrophobic contacts with the ribose rings of the two recognized cytosines (Fig 4). However, a mutation of the phenylalanine to a leucine, another hydrophobic residue, might not be sufficient to prevent the binding of RRM1 to RNA as it may still allow these hydrophobic contacts. This could explain the absence of effect of this protein mutant on Npl3 functions (Colombo et al., 2017, Lee-Soety et al., 2012). Our structural-guided analysis uncovered the functional contribution of RRM1 binding to RNA *in vivo*.

In humans, SR proteins were previously shown to recruit U1 and U2 snRNPs on the 5’SS and 3’SS, respectively (Long & Caceres, 2009). In yeast, the use of a *npl3 Δ* strain revealed a general decrease of pre-mRNA splicing suggesting a role in constitutive splicing. Npl3 was reported to facilitate co-transcriptional splicing by recruiting the U1 snRNP on RNA pol II (Kress et al., 2008). In addition, the protein was recently shown to interact with U1 snRNP through protein-protein interaction using its RGG/RS domain (Muddukrishna, Jackson et al., 2017). However, we could not detect in our iCRAC data any specific enrichment of Npl3 binding around spliced introns nor specific enrichment in ribosomal protein genes. In good agreement with this observation, it was previously proposed that the recruitment of Npl3 at these sites might be driven through interactions with other proteins and chromatin modifications (Kress et al., 2008, Moehle et al., 2012). Our split-iCRAC data show a specific binding of Npl3 to the Stem-loop I of the U2 snRNA. Moreover, RNA mutations that prevent the binding of Npl3 without affecting the stability of the Stem-loop I resulted in a slow growth phenotype in combination with the deletion of *lea1*, as observed with Npl3 mutants preventing its binding to RNA (Fig. 6). We found that SRSF1 (Jobbins, Campagne et al., 2022) and FUS (Jutzi, Campagne et al., 2020) could interact directly with the SL3 of the human U1 snRNA. The specific interaction between Npl3 and the U2 snRNA reported here implicates a broader role for the U snRNAs. For example, this interaction could serve as an early and transient binding platform to load splicing factors having a specific function during the splicing reaction. In addition, this interaction of Npl3 with U2 snRNA stem-loop I suggests an unexpected mode of action of this protein in splicing. As Npl3 was shown to genetically interact with Snu66, a component of the tri-snRNP (Kress et al., 2008), its binding to U2 snRNA could play a role at a later stage of the spliceosome assembly. Our structure shows that Npl3 can unfold the stem-loop I by binding to its RNA target sequence. This unfolding is required for the recruitment of the U4/U6.U5 tri-snRNP as this unfolded sequence can then form a duplex with the U6 snRNA upon the spliceosome complex A to B transition, which leads to the active B^act^ spliceosome complex (Sashital et al., 2007, Yan et al., 2016). Therefore, Npl3 may play an early chaperoning role for U2-U6 hybridization, which would facilitate the formation of the B^act^ complex. This would also explain why the effect of the stem-loop I mutation on yeast growth is more visible at low temperature, as the dynamics of the RNA rearrangement may be slower and the role of Npl3 more important to unfold U2 stem-loop I. In agreement with its involvement at this stage of the splicing reaction, Npl3 was proposed to stimulate Prp28’s ATPase activity to remove U1 snRNP from the pre-B complex (Yeh et al., 2021). Moreover, Npl3 was detected by mass spectrometry in the B complex (Fabrizio, Dannenberg et al., 2009). Interestingly, 16 nts of the U2 snRNA encompassing the sequence targeted by Npl3 were not visible in the cryo-EM structure of the yeast B complex (Bai, Wan et al., 2018) indicating some flexibility near the U2-U6 helix II duplex formation. In addition, a large empty cavity is present at this location which is large enough to perfectly accommodate the two Npl3 RRMs bound to RNA (Fig. S12). Moreover, the α-helix from an unidentified protein was observed at the proximity of this Npl3 binding site in the cryo-EM structure of the yeast spliceosome complex C (Galej, Wilkinson et al., 2016). All these biochemical and structural elements point towards an RNA chaperoning activity of Npl3 in the formation of active spliceosomes in yeast and pave the way for mechanistic investigation on the mode of action of other SR- and SR-like proteins in higher eukaryotes.

## Materials and Methods

### Expression and purification of recombinant protein

Escherichia coli BL21 (DE3) codon plus cells transformed with pET28a::Npl3 RRM1 (residues 114-201), pET28a::Npl3 RRM2 (residues 193-282) or pET28a::Npl3 RRM1/2 (residues 114-282) were grown at 37 °C in M9 minimal medium supplemented with 50 μg/ml kanamycin, 34 μg/ml chloramphenicol, 1 g/l ^15^NH_4_ Cl and 4 g/l unlabeled or 2 g/l ^13^C labeled glucose for ^15^N or ^15^N and ^13^C labeled proteins, respectively. Protein expression was induced at OD_600_ of 0.9 with 1 mM IPTG at 20 °C. After 18 hours, the cells were harvested, and proteins were purified by two successive nickel affinity chromatography (Qiagen_®_) steps. The proteins were dialyzed in RRM1 NMR buffer (25 mM Na_2_HPO_4_, 25 mM NaH_2_PO_4_, pH 6.9), RRM2 NMR buffer (100 mM NaCl, 20 mM NaH_2_PO_4_ pH 5.5), or RRM1/2 NMR buffer (25 mM Na_2_HPO_4_, 25 mM NaH_2_PO_4_, pH 6.9). The concentration of recombinant proteins was carried out using 10-kDa molecular mass cut-off Centricons (Vivascience_®_). The absence of RNases was confirmed using the RNase Alert Lab Kit (Ambion_®_).

### Preparation of RNA–protein complexes

All RNA oligonucleotides were purchased from Dharmacon_®_, de-protected according to manufacturer’s instructions, lyophilized and resuspended in the corresponding NMR buffer. NMR titrations were carried at a protein concentration of 0.2 mM. The Npl3 RRM–RNA complexes used for structure calculations were prepared in their corresponding NMR buffer at a protein:RNA stoichiometric ratio of 1:1 and a final concentration of 0.9 mM. The U2 RNA stem-loop (5’-GAGCGAAUCUCUUUGCCUUUUGGCUUAGAUC-3’) was transcribed *in vitro* and purified by HPLC on an anion exchange column at 85°C and in denaturing conditions (6 M urea). The fraction containing the RNA was precipitated using butanol and dissolved in water. The RNA was renaturated 30 sec at 95°C followed by a slow cooling step to room temperature.

### Isothermal titration calorimetry

ITC experiments were performed on a VP-ITC instrument (Microcal), calibrated according to the manufacturer’s instructions. Protein and RNA samples were dialyzed against the NMR buffer. Concentrations of proteins and RNAs were determined using optical-density absorbance at 280 and 260 nm, respectively. 10 μM of each RNAs were titrated with 200 μM of protein by 40 injections of 6 μl every 5 min at 40°C. Raw data were integrated, normalized for the molar concentration and analyzed using the Origin 7.0 software according to a 1:1 RNA:protein ratio binding model.

### NMR experiments

All the NMR spectra were recorded at 313 K using Bruker AVIII-500 MHz, 600 MHz, 700 MHz, AVIIIHD-600 MHz, 900 MHz equipped with a cryoprobe, and AVIII-750 MHz spectrometers. Topspin 3.0 (Bruker_®_) was used for data processing and Sparky (http://www.cgl.ucsf.edu/home/sparky/) for data analysis.

Protein backbone assignment was achieved using 2D ^1^H–^15^N HSQC and 3D HNCACB, while side chain assignments were achieved using 2D ^1^H–^13^C HSQC, 3D HcccoNH TOCSY, 3D hCccoNH TOCSY, 3D NOESY ^1^H–^15^N HSQC and 3D NOESY ^1^H–^13^C HSQC aliphatic. Aromatic protons were assigned using 2D ^1^H–^1^H TOCSY and 3D NOESY ^1^H–^13^C HSQC aromatic (Sattler, 1999).

RNA resonance assignments in complex with Npl3 RRMs were performed using 2D ^1^H–^1^H TOCSY, natural abundance 2D ^1^H–^13^C HSQC and 2D ^13^C 1F-filtered 2F-filtered NOESY in 100% D_2_O. Intermolecular NOEs were obtained using 2D ^1^H–^1^H NOESY and 2D ^13^C 2F-filtered NOESY (Peterson, Theimer et al., 2004) in the presence of unlabeled RNA and ^15^N- and ^15^N-^13^C-labeled proteins, respectively.

All NOESY spectra were recorded with a mixing time of 100 ms, the 3D TOCSY spectrum with a mixing time of 17.75 ms and the 2D TOCSY with a mixing time of 60 ms.

The 1D NMR experiments shown in Fig. 6B were recorded at 298 K on a Bruker 700 MHz spectrometer in the RRM1/2 NMR buffer at RNA concentrations of 50 μM.

^15^N T1 and T2 measurements were recorded at 313 K at a ^1^H frequency of 600 MHz with established methods as previously described (Barraud & Allain, 2013).

### Residual dipolar couplings (RDCs) measurement

In order to measure the amide residual dipolar couplings of Npl3 RRM1/2 in its RNA bound state, we used Pf1 phages as alignment medium. The Pf1 phages were previously washed using the NMR buffer according to the manufacturer instructions (Alsa Biotech). The Pf1 phages were then added to the sample at a concentration of 10 mg/ml and the formation of a crystalline medium was monitored by measuring the splitting of the deuterium atoms from D_2_O. In order to extract amide RDCs, we compared the apparent scalar couplings H-N observed on a 2D ^1^H-^15^N IPAP HSQC before and after addition of the phages. The RDC restraints were then added during the cartesian refinement procedure using AMBER.

### Structure calculations

AtnosCandid software (Herrmann, Guntert et al., 2002) was used for peak picking 3D NOESY (15 N- and 13 C-edited) spectra. Preliminary structures and a list of automatically assigned NOE distance constraints were generated through 7 cycles using CYANA *noeassign* (Herrmann et al., 2002). Additionally, intra-protein hydrogen bond constraints were added based on hydrogen–deuterium exchange experiments on the amide protons. For these hydrogen bonds, the oxygen acceptors were identified based on preliminary structures calculated without hydrogen bond constraints. Intra-molecular RNA and RNA–protein intermolecular distance restraints were manually assigned and added to the calculation with 62 RDCs restraints in the case of Npl3 RRM1/2. Calculations with the RNA were done using CYANA 3.98.4 in which seven iterations were performed, and 500 independent preliminary structures were calculated at each iteration step. These 50 structures were refined with the SANDER module of AMBER 14.0 (Case, Cheatham et al., 2005) by simulated annealing in implicit water using the rna.ff12SB force field (Lindorff-Larsen, Piana et al., 2010). The 10 best structures based on energy and NOE violations were analyzed with PROCHECK (Altvater, Chang et al., 2012, Janke, Magiera et al., 2004, Laskowski, Rullmannn et al., 1996, Longtine, McKenzie et al., 1998). Figures were generated with MOLMOL (Koradi, Billeter et al., 1996) and Pymol. The Ramachandran plot of the Npl3 RRM1 in complex with RNA indicates that 86.9% of the residues are in the most favored regions, 12.9% in the additional allowed regions, 0.1% in the generously allowed regions and 0% in the disallowed regions. The Ramachandran plot of the Npl3 RRM1/2 in complex with RNA indicates that 71.2% of the residues are in the most favored regions, 25.4% in the additional allowed regions, 3.3% in the generously allowed regions and 0.1% in the disallowed regions.

## Modified Scaffold independent analysis

The method was adapted from Beuth et al. 2007 (Beuth et al., 2007). Briefly, ^1^H–^15^N HSQC NMR titrations were done with 0.2 mM Npl3 RRM1 protein in the RRM1/2 buffer at 30 °C with successive addition of ssDNA (RNA:protein ratio 0.3:1, 0.6:1, 1:1, 2:1). The chemical shift perturbations observed were calculated for the 1:1 ratio with the formula (Δδ = [(δHN)^2^ + (δN/6.51)^2^]^1/2^). The values calculated for non-overlapping peaks were summed.

### Split-iCRAC

The BY4741 (MATa ura3 his3 leu2 met15 TRP1) was used as a parental yeast strain in which the promoter of Npl3 was replaced with an inducible gal promoter by homologues recombination. The strain was complemented with pRS315::Npl3±500kbp::HRV-3C::CterHTP plasmid. In this plasmid, the expression of Npl3 is driven by its endogenous promoter. A HRV-C3 protease cleavage site was inserted between amino acids S282 and N283 using PCR. A His-Trypsin-Protein A (HTP) tandem tag was placed at the C-terminus of the protein by PCR.

The split-iCRAC was based on the original CRAC protocol described by Granneman et al. 2009 with some modifications (Granneman, Kudla et al., 2009). Briefly, the recombinant yeast strains were grown in SD-leu medium to drive Npl3 expression only from the transformed plasmid. 2 L of yeast culture were harvested at an OD600 of ∼2. The cells were resuspended in 1 v/w SD-Trp medium and UV-irradiated (1.6 J/cm^2^) in Petri dishes in a Stratalinker 1600 (Stratagene). Half the cells were not subjected to UV treatment and were kept as the UV minus control. Cells were pelleted and resuspended in 25 ml of lysis buffer (50 mM Tris-HCl pH 7.8, 150 mM NaCl, 0.15% NP-40) with addition of 1.3 mM PMSF, 1 mM DDT, complete protease inhibitor cocktail (Roche_®_)). The cells were lysed using 25 ml glass beads in Planetary mill for 20 mins at 500 rpm. 5 ml of lysis buffer was added, and the lysate was cleared by centrifugation (20 min at 5000 rpm and 4 °C followed by 2x 20 min at 18000 rpm and 4 °C). Cleared lysate was incubated with 300 of IgG bead suspension (1:1) for 2 h at 4 °C with rotation. Beads were collected and washed 3x with 10 ml wash buffer (50 mM Tris-HCl pH 7.8, 1 M NaCl, 0.15% NP-40, 0.5 mM DDT) followed by 3x with 10 ml lysis buffer with added 0.5 mM DDT. Beads were resuspended in 5 ml of lysis buffer and rotated with 40 μg of homemade TEV for 18 h at 4 °C. The eluates were concentrated to a volume of 500 μl in 30 kDa cutoff centricons (Millipore_®_). No RNase treatment was performed in the final experiments for library preparation. 0.4 g of Guanidine-HCl were dissolved in the eluate to yield a final concentration of 6M. NaCl and imidazole were added to a final concentration of 300 and 10 mM, respectively. 100 μl of pre-equilibrated Ni-beads were added to the samples and incubated for 2h at 4 °C. Beads were washed 3x with wash buffer (6 M guanidine-HCl, 50 mM Tris-HCl (pH 7.8), 300 mM NaCl, 0.1% NP-40, 40 mM imidazole, and 1 mM DDT) followed by 3x with PNK buffer (50 mM Tris-HCl (pH 7.8), 40 mM MgCl_2_, 0.1% NP-40, and 1 mM DDT). The beads were then incubated in 50 μl of PNK buffer (pH 6.5) in the presence of 10 units of PNK (NEB_®_) and 10 units of CIP (NEB_®_) and 20 units of SUPERase.In (Ambion_®_) for 30 mins at 37 °C. Beads were washed 3x with wash buffer followed by 2x with PNK buffer (pH 7.8) and incubated with 50 pmoles of L3 adapter (rAppAGATCGGAAGAGCGGRRCAG/ddC/) in 50 μl of ligation mixture (1x PNK buffer (pH 7.8) supplemented with 12.5 units of T4 RNA ligase 1 (NEB_®_), 250 units of T4 RNA ligase 2 truncated K227Q (NEB_®_), 40 units of SUPERase.In (Ambion_®_) and 10% PEG 4000). The reaction is incubated for 18 h with mild shaking at 16 °C. Beads are then washed with 3x wash buffer followed by 3x PNK buffer. For the PreScission cleaved version, the beads are then incubated with 1x PNK buffer (pH 7.8) supplemented with 20 units of SUPERase.In (Ambion_®_) and 10 units of PreScission protease (GE Healthcare_®_) for 18 h at 4 °C with mild shaking. The mixture is then supplemented with 10 units of PNK and 0.5 μl of γ^32^P-ATP and incubated for 1 h at 37 °C. 20 μl of 4x LDS is added to the beads, boiled for 10 mins and resolved on a 4-12% Bis-Tris NuPage gel (Invitrogen_®_). Subsequent steps of RNA isolation and library preparation were done as described in Huppertz et al. 2014 Methods (Huppertz et al., 2014). The negative control sample was generated following the same procedure but starting with WT yeast strain that does not express any tagged protein.

### High-throughput sequencing and analysis

Four replicates of the full-length protein and RRM1/2 domain, three replicates of the RGG/RS and one negative control sample were Illumina sequenced on a single lane of the NextSeq500 High Output (single-end 75 bp reads) according to manufacturer’s protocol. The reads corresponding to each sample were demultiplexed using the FLEXBAR tool (Dodt, Roehr et al., 2012). At least 25 million reads were generated (up to 50 million) for each sample. After barcode removal and quality trimming, the reads were mapped against the S. cerevisiae reference genome S288C_R64-1-1 (Engel, Dietrich et al., 2014) using STAR (Dobin, Davis et al., 2013). Approximately 30% of the reads of each sample were uniquely mapped to the genome. Reads mapping to the same genomic location and with the same Unique Molecular Identifier (UMI) were assumed to arise from PCR duplication and were therefore merged using UMI-tools (Smith, Heger et al., 2017). To increase the reliability of identifying significant crosslinking sites, the deduplicated reads from the replicate samples were merged and subsequently used for peak calling using iCount (Sugimoto, Konig et al., 2012). A False Discovery Rate (FDR) cut-off of 0.05 was used to identify significant crosslinking sites. Identified crosslinking sites that were less than three nucleotides apart were merged into one cluster. Clusters that overlapped with ones identified in the negative control sample were not included in subsequent analysis steps. HOMER software (Heinz et al., 2010) was used for de novo motif discovery using the identified clusters flanked by five nucleotides on each side because of the expected short motif. Motif predictions were made for sizes ranging from 2 up to 10 nucleotides. The HOMER motifs were validated by plotting the density of the top motifs in the different samples and the negative control in a window of ±50 nucleotides around the crosslink cluster centers.

### Mutant growth analysis

Genetically modified yeast strains were prepared by homologues recombination according to standard protocols (Altvater et al., 2012, Janke et al., 2004, Longtine et al., 1998). Equal amounts of streaked yeast cells were resuspended in 90 μl H_2_O and subsequently diluted 10x in a series of 5 steps. 10 μl of the four lowest dilutions were spotted on SD plates. Plates were incubated up to one week at appropriate temperatures and pictures were taken daily using a Coolpix P310 digital camera (Nikon^®^).

### Data deposition

The coordinates of the Npl3 RRM1–AUCCAA (PDB ID 7QDD) and Npl3 RRM1/2-AUCCAGUGGAA (PDB ID 7QDE) structures have been deposited in the Protein Data Bank (PDB),www.pdb.org. The split-iCRAC data are available at http://www.ebi.ac.uk/arrayexpress/ using this ArrayExpress accession number E-MTAB-11736.

## Acknowledgements

We thank Dr. Alvar Gossert, Dr. Simon Rudisser and Dr. Fred Damberger for the maintenance of the biomolecular NMR spectroscopy platform. We also thank the SNF (grant numbers: 310030B-189379 and 31003A-170130) and SNF-NCCR RNA and Disease (grant number: 51NF40-182880). V. G. Panse acknowledges financial support from the Swiss National Science Foundation, NCCR RNA & Disease, Novartis Foundation, Olga Mayenfisch Stiftung and a Starting Grant Award (EURIBIO260676) from the European Research Council.

## Author contributions

FHTA, VGP and AC designed research. SG, SR, AM and AC prepared samples. AM, SG, SR and AC collected data. AM, AC, SG, KMB, SR, SC, IB and MMD analyzed data. AM and AC wrote the manuscript and all authors edited it.

## Conflict of interest

The authors declare that they have no conflict of interest.

## Figure Legends

**Supplementary figure 1: Top enriched motifs of the HOMER de novo motif search for the split-iCRAC. (A)** The top 4 enriched HOMER motifs in the crosslink clusters ±5 nt for the different constructs of Npl3 compared to the random background. The p-value of each motif indicates its enrichment in the corresponding sample over random sequences of similar length from the S. cerevisiae genome. **(B)** Density plot for each of the enriched motifs in the 100 nucleotides around the crosslinking cluster centers in the different samples and the negative control. Traces for the full-length protein, RRM1/2, RS and negative control are colored in purple, blue, green and yellow, respectively. All motifs show similar densities in the different samples compared to the negative control except the three motifs boxed in red. Those three motifs have significantly higher densities immediately preceding the crosslinking cluster center in the FL and RRM1/2 samples.

**Supplementary figure 2: Representation of the correlation time (τ**_**c**_**) measured in the presence of Npl3 RRM1/2 bound to the AUCCAGUGGAA RNA**. The two RRMs tumbled with RNA as a single unit.

**Supplementary figure 3: Comparison of the binding of Npl3 RRM1/2 to the 5’-AUCCAGUGGAA-3’ RNA with isolated RRM complexes**. Representation of the combined chemical shift perturbations of Npl3 RRM1/2 amide residues upon binding to the 5’-AUCCAGUGGAA-3’ RNA (in blue) at a ratio of 1:1 in comparison with the isolated RRM domains bound to their respective RNA targets (in red). The corresponding secondary structure elements are represented at the top of the graph. RRM1, RRM2 and the inter-domain linker are colored in gray, green and magenta, respectively. Some residues could not be assigned in the bound form of RRM1/2 due to the intermediate to slow exchange regime.

**Supplementary figure 4: Npl3 RRM1 binds preferentially to polyC sequences**. Overlay of ^1^H-^15^N HSQC spectra recorded with Npl3 RRM1 upon titration with 6mer polyA, polyG, polyT and polyC ssDNA as well as 8mer polyU RNA sequences. The spectra were recorded at 30°C, in the Npl3 RRM1/2 NMR buffer. Blue represents the free protein while red represents the bound protein at a 1:1 ratio.

**Supplementary figure 5: Study of the interaction of Npl3 RRM1/2 with an RNA containing inverted RRM binding sites (AUGGAGUCCAA). (A)** NMR overlay of ^1^H-^15^N HSQC spectra recorded with Npl3 RRM1/2 free form, bound to AUGGAGUCCAA at 1:0.3 and 1:1 ratios and to AUCCAGUGGAA at a 1:1 ratio. **(B)** ITC measurement performed with Npl3 RRM1/2 and the AUGGAGUCCAA RNA.

**Supplementary figure 6: ITC measurements performed with the AUCCAGUGGAA RNA and different variants of the Npl3 RRM1/2 protein**.

**Supplementary figure 7: ITC measurements performed with Npl3 RRM1/2 and different variants of the AUCCAGUGGAA RNA**.

**Supplementary figure 8: Point mutations in RRM1 and RRM2 disrupt their binding to nucleic acid without affecting the fold of the domains**. (**A**) Overlay of ^1^H-^15^N HSQC spectra recorded with RRM2 mutants in their free form and upon addition of their target sequence at a 1:1 ratio represented in blue and red, respectively. The spectra were recorded at 40°C, in the RRM2 NMR buffer. Binding was tested to 5’-ATGGTC-3’ ssDNA except for the Q214A, K217A double mutant which was done with 5’-AUGGUC-3’ RNA. The mutated residues are highlighted on the structure of the RRM2-RNA complex (**B**) Overlay of ^1^H-^15^N HSQC spectra recorded with RRM1 mutants in their free form and upon addition of their target sequence at a 1:1 ratio represented in blue and red, respectively. The spectra were recorded at 40°C, in the RRM1 NMR buffer. Binding was tested to 6mer polyC ssDNA. The mutated residues are highlighted on the structure of the RRM1-RNA complex. The spectrum of the double mutant F128A, F160A in the bound form was not recorded, but ITC measurement indicates a strong decrease in affinity of this mutant for RNA (Fig. S6).

**Supplementary figure 9: Direct binding of Npl3 to the U2 snRNA and not the U1 snRNA**.Genome browser view of the LSR1 gene encoding the U2 snRNA displaying the split-iCRAC coverage tracks of the full-length protein, RRM1/2, RGG/RS constructs and the negative control. Blue boxes represent the identified crosslinking clusters. The enlarged sequence represents the sequence of the 5’ region of the gene that is represented in the crosslinking clusters of the three protein constructs. **(B)** Similar view for the snR19 gene encoding the U1 snRNA showing that no specific crosslinking cluster could be detected with Npl3.

**Supplementary figure 10: (A)** Barplot of the read distribution across different RNA species in the split-iCRAC experiment. Each bar represents one sample and the y-axis the percentage of unique reads mapping to different RNA species as annotated in the Ensembl reference annotation of S. cerevisiae (Saccharomyces_cerevisiae.R64-1-1.83). **(B)** The distance between the RS/RGG domain and the closest RRM cluster midpoint is shown. The RS domain binds RNA on both sites of RRMs crosslinking sites.

**Supplementary figure 11: Comparison of the mode of interaction of Npl3 and other RNA binding proteins with RNA. (A)** Structures of Npl3 RRM1 and SRSF2 RRM bound to RNA. The color code is as in Fig. 4A. The two RRMs bind similarly to two consecutive cytosines.**(B)** Schematic representation of the binding interaction of Npl3, TDP43 and Dnd1 RRM1/2 with RNA. The RNA is represented by a black arrow, the inter-domain linker is in red and the part of RRM2 involved in the interaction is colored in blue, yellow and cyan, respectively.

**Supplementary figure 12: Npl3 RRM1/2 bound to RNA fits into a cavity present at its U2 snRNA cross-linked site in the yeast spliceosome complex B. (A)** View of the cryo-EM structure determined with the yeast spliceosome complex B. A large cavity is observed at the position where the Npl3 cross-linked sequence is not visible. **(B)** The structure of Npl3 RRM1/2 bound to RNA fits perfectly in this cavity.

## Notes

### Competing Interest Statement

The authors have declared no competing interest.

