## Supplementary figures and images for "The structure of yeast Npl3 bound to RNA reveals a cooperative sequence-specific recognition and an RNA chaperone role in splicing"

### Supplemental Figure 1

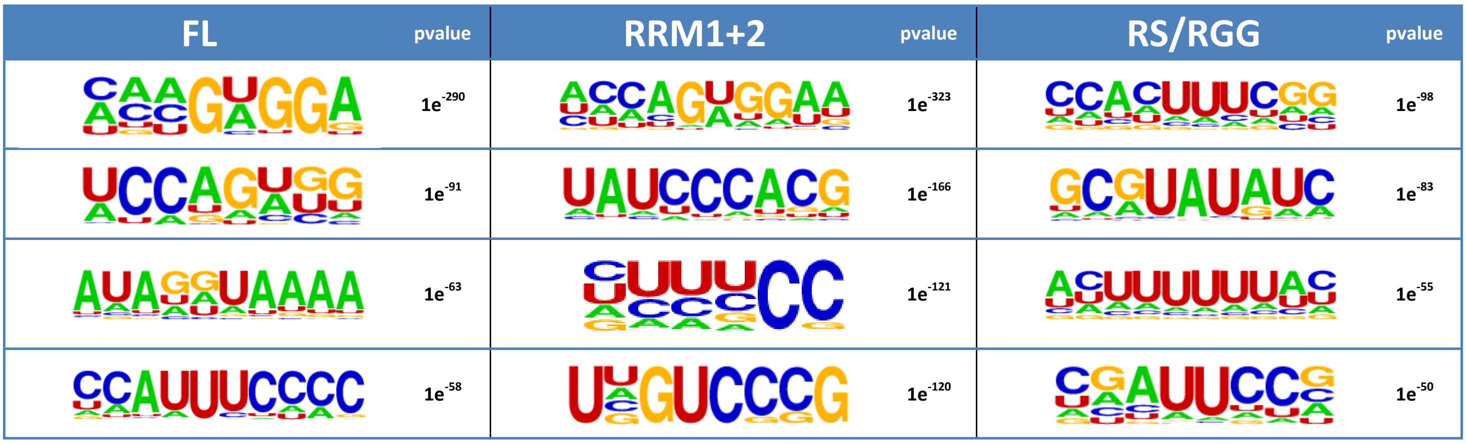

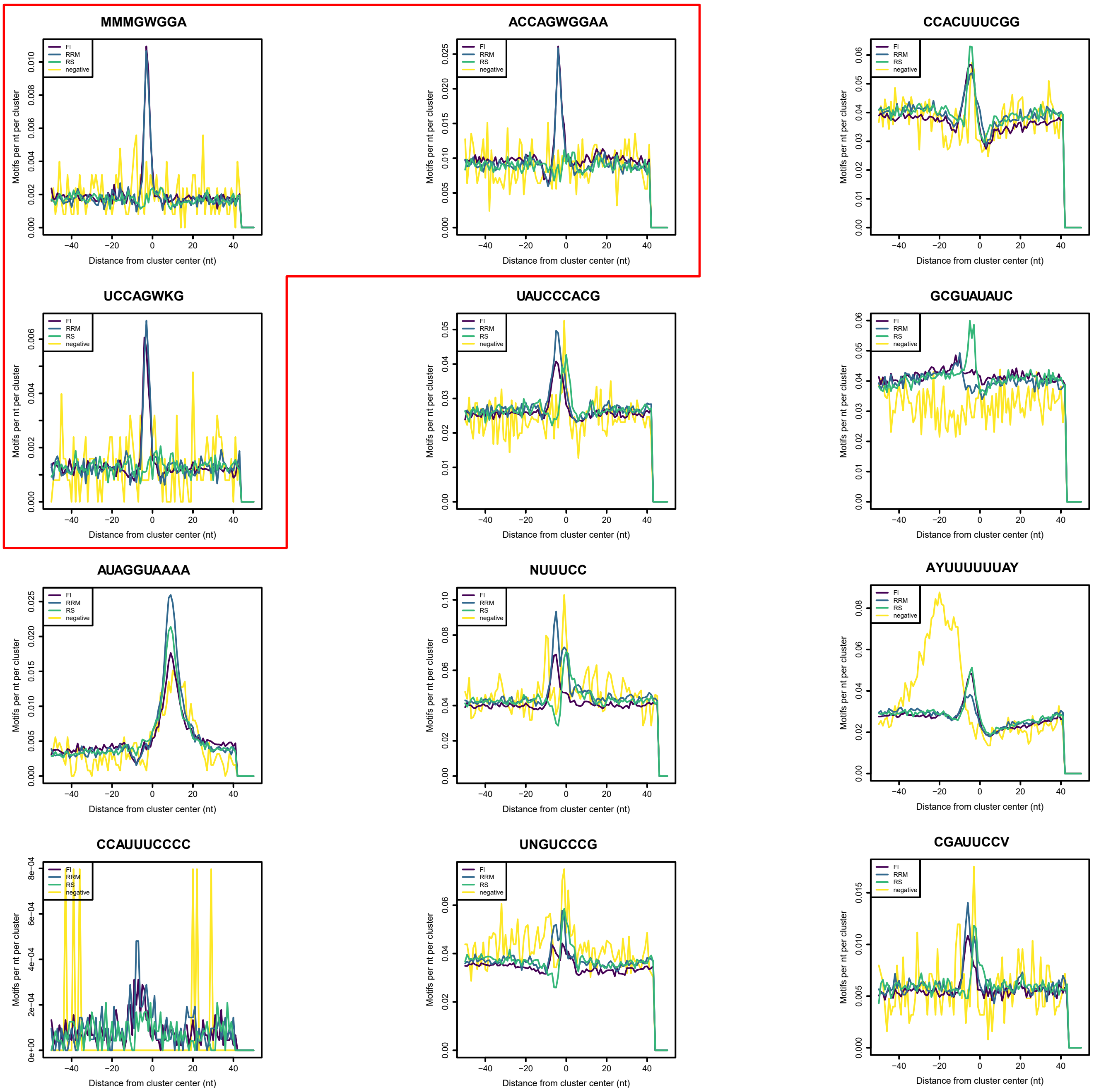

### Supplemental Figure 2

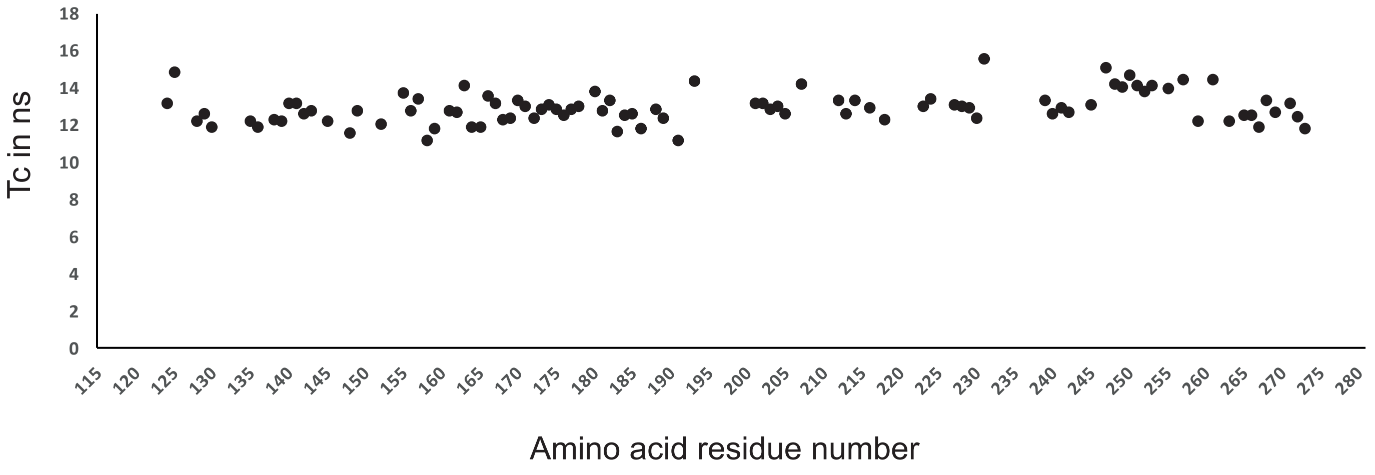

### Supplemental Figure 3

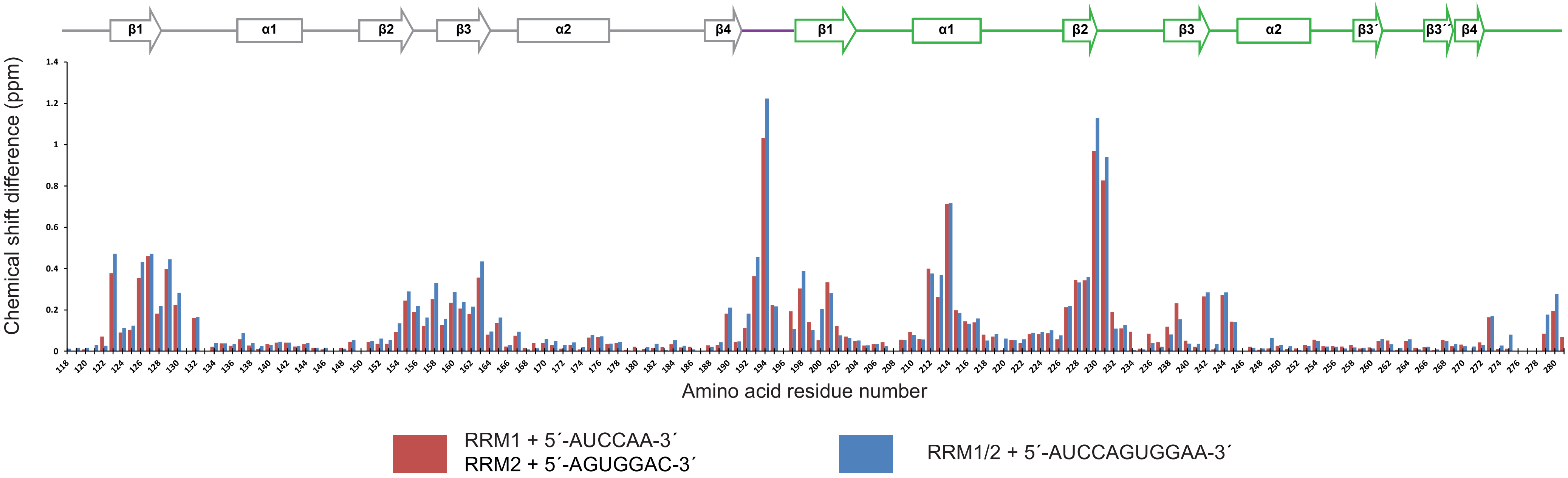

### Supplemental Figure 4

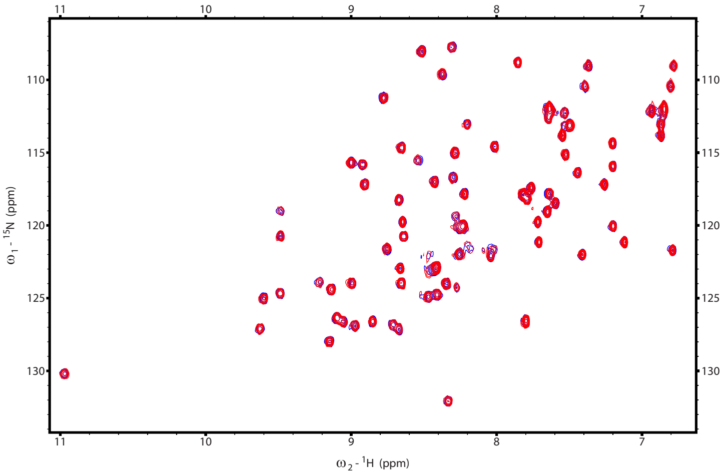


PolyA

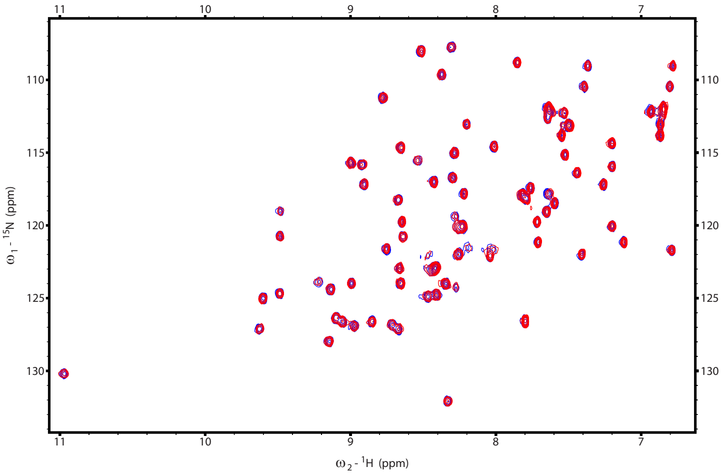


PolyG

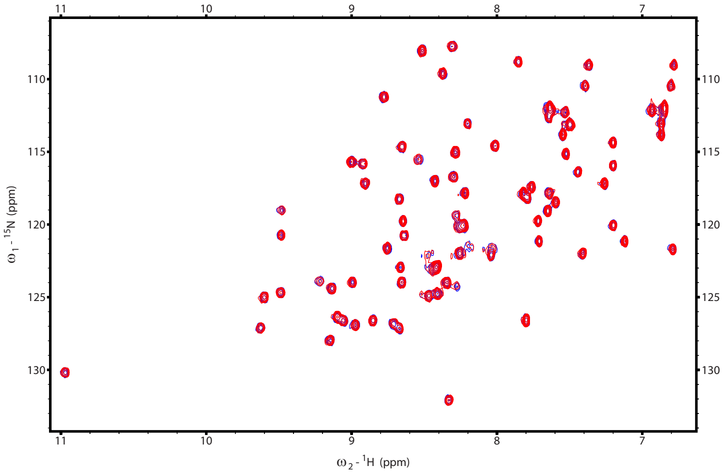


PolyT

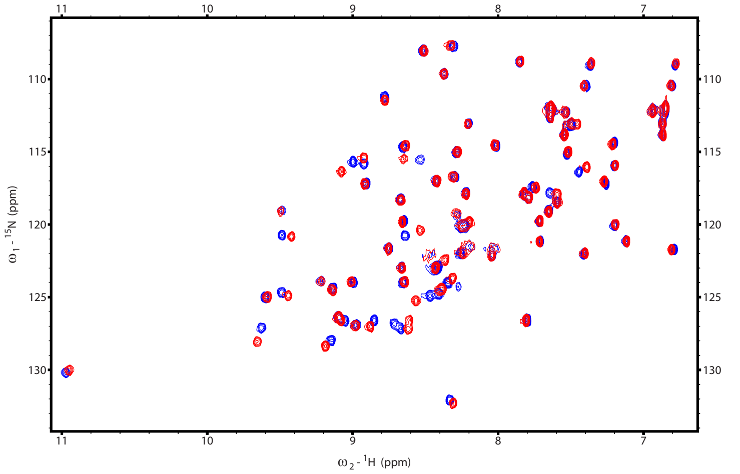


PolyC

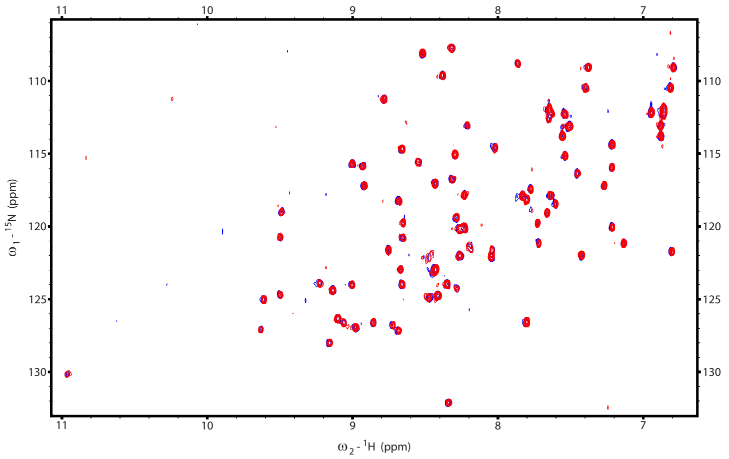


PolyU

Figure S4

### Supplemental Figure 5

A

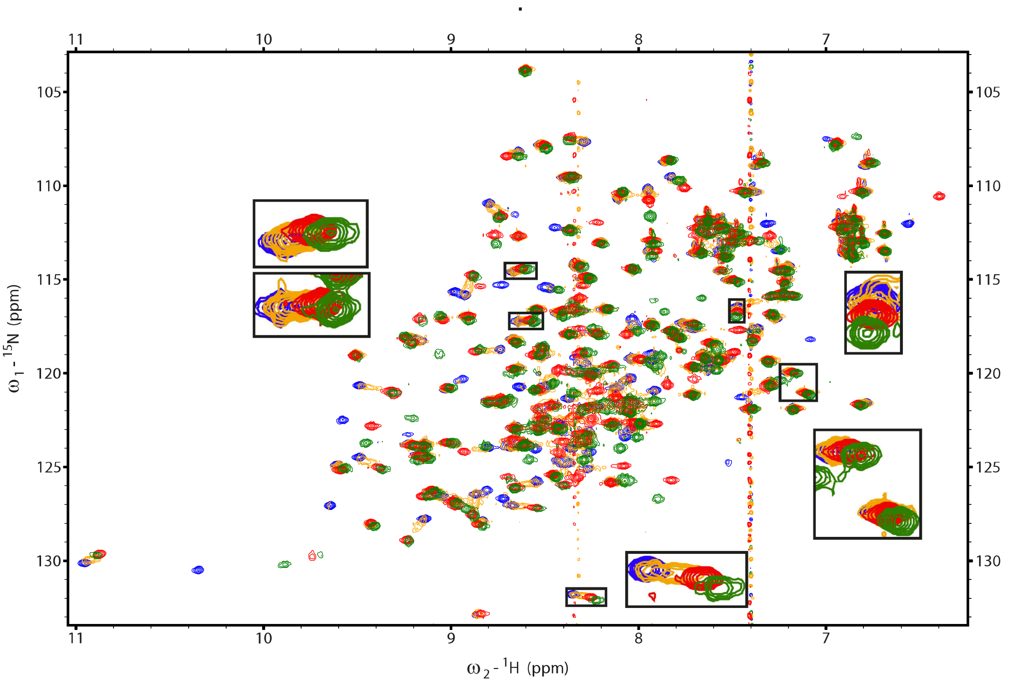


RRM1/2 free   RRM1/2:AUGGAGUCCAA 1:0.3   RRM1/2:AUGGAGUCCAA 1:1   RRM1/2:AUCCAGUGGAA 1:1

B

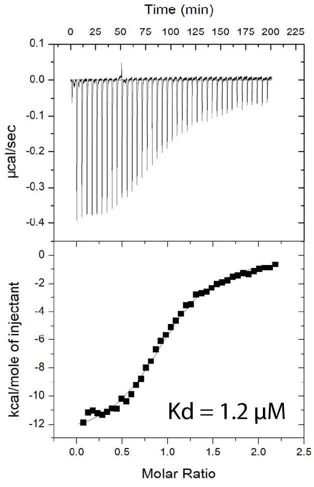


Figure S5

### Supplemental Figure 8

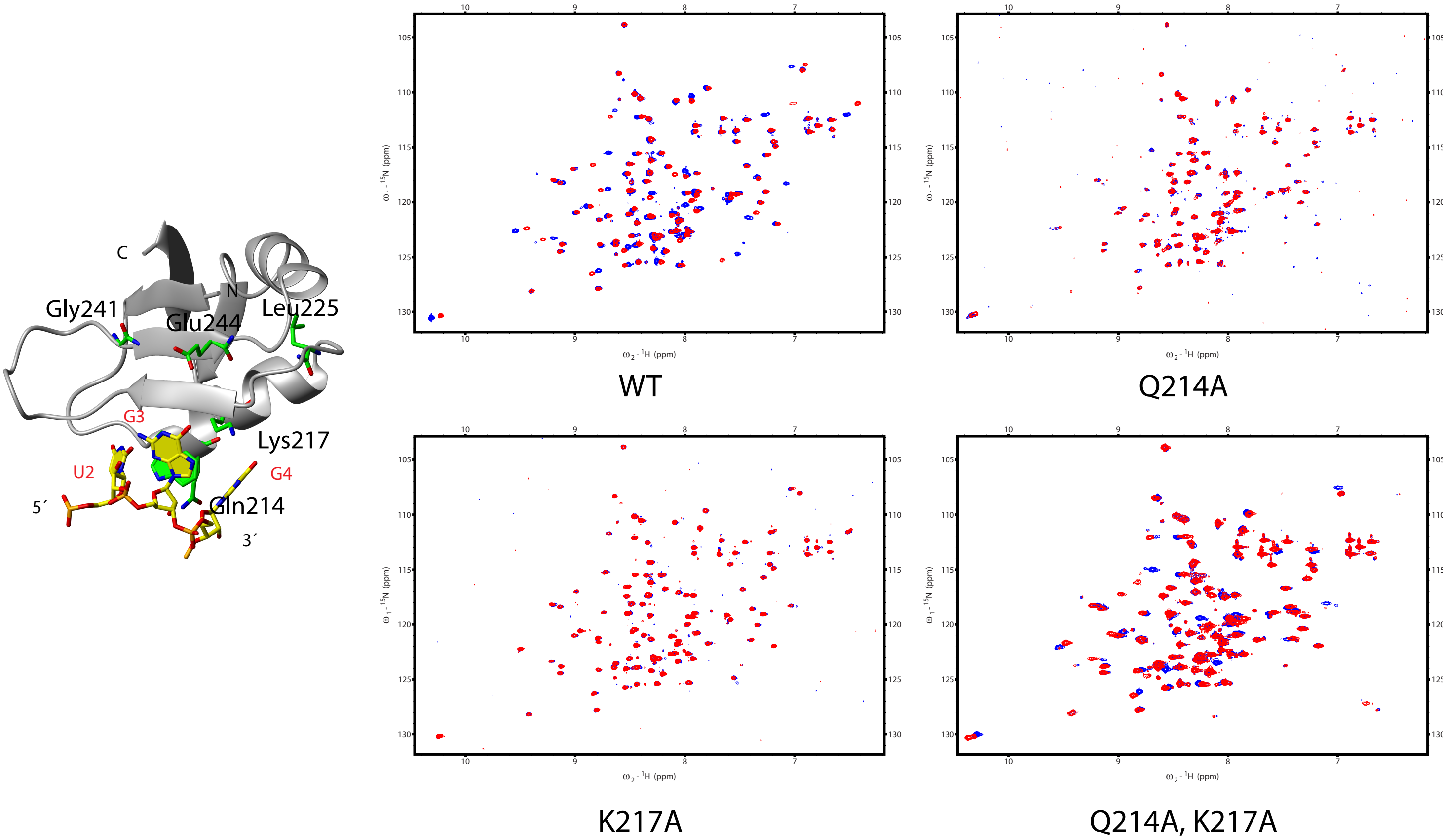

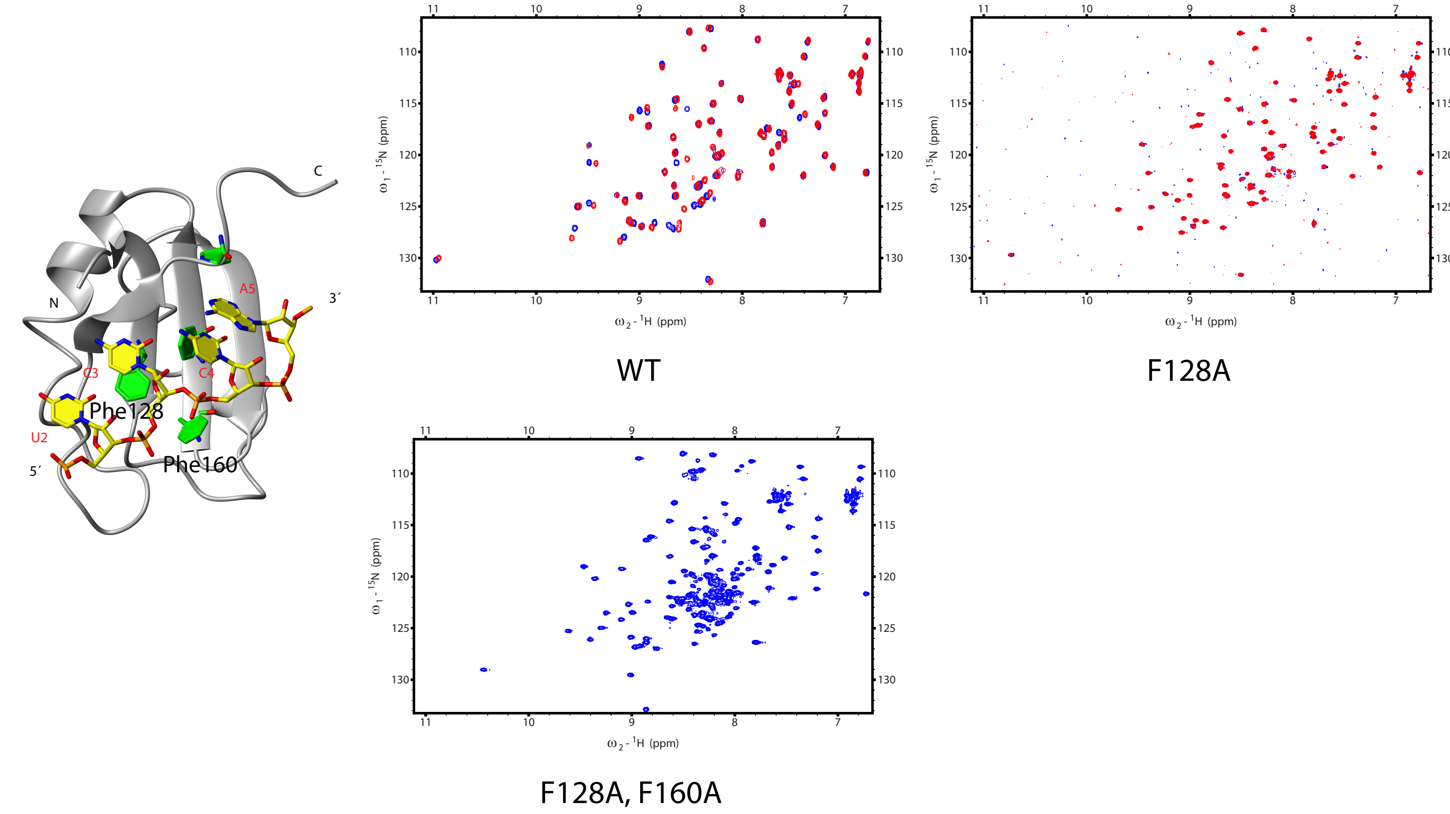

### Supplemental Figure 9

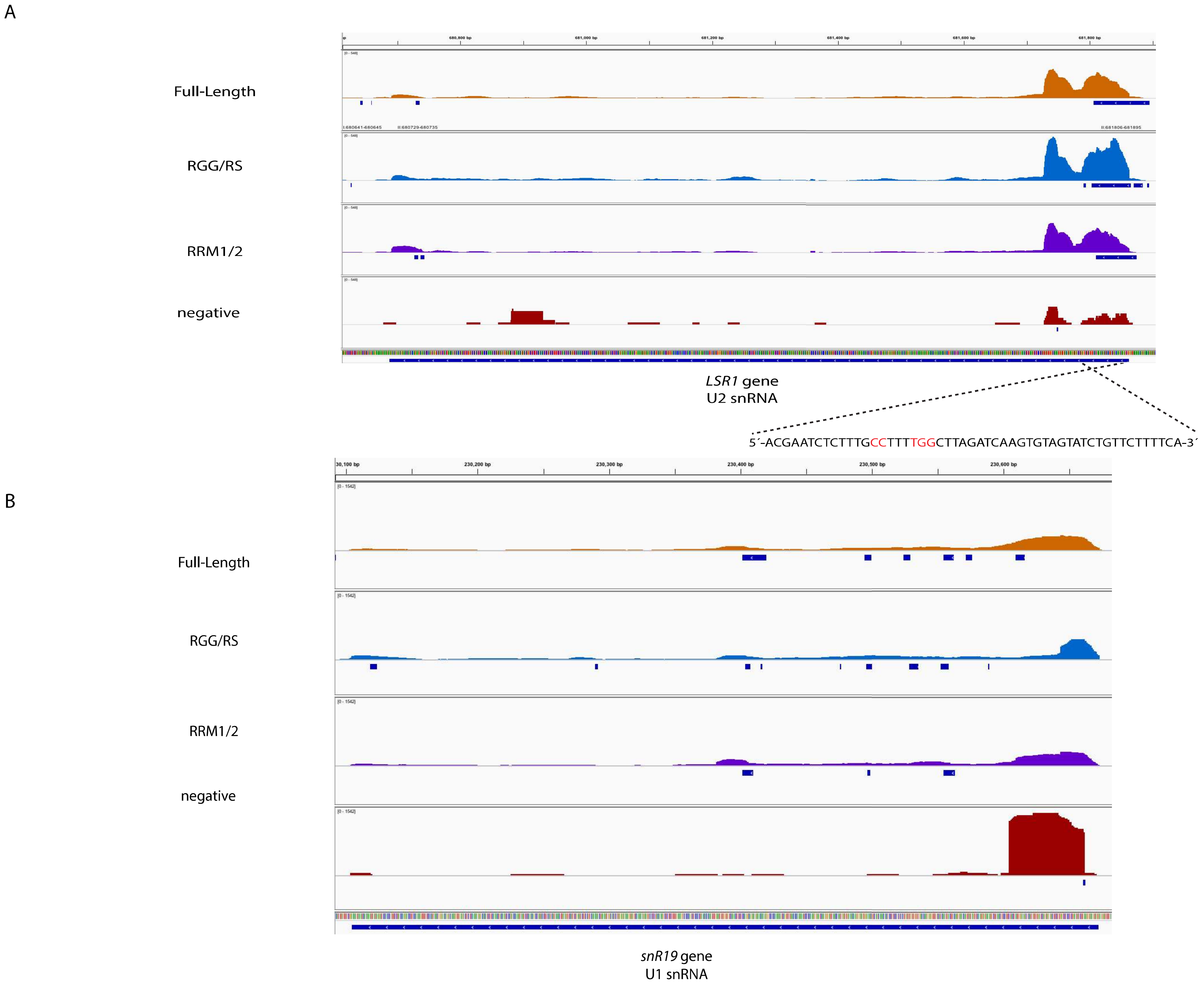

### Supplemental Figure 10

A

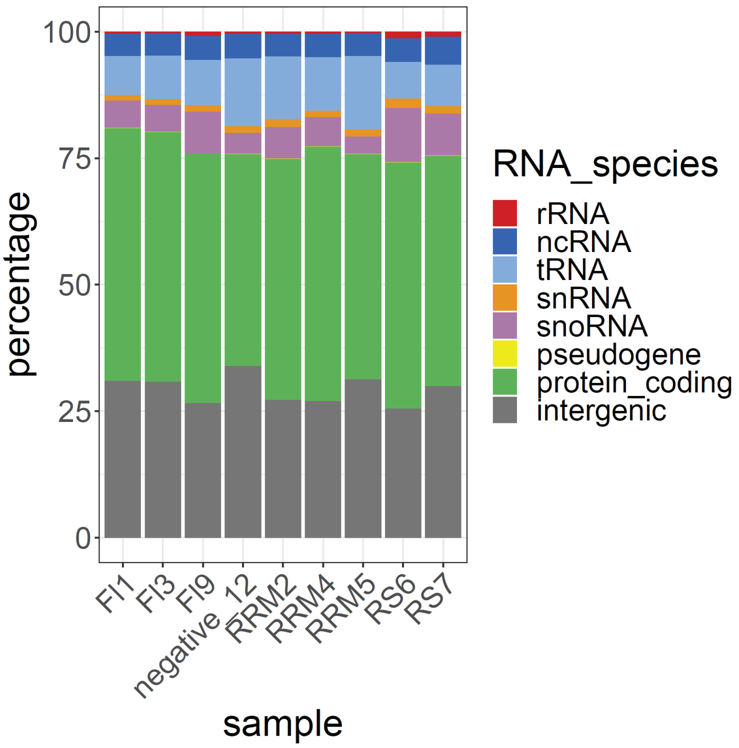


B

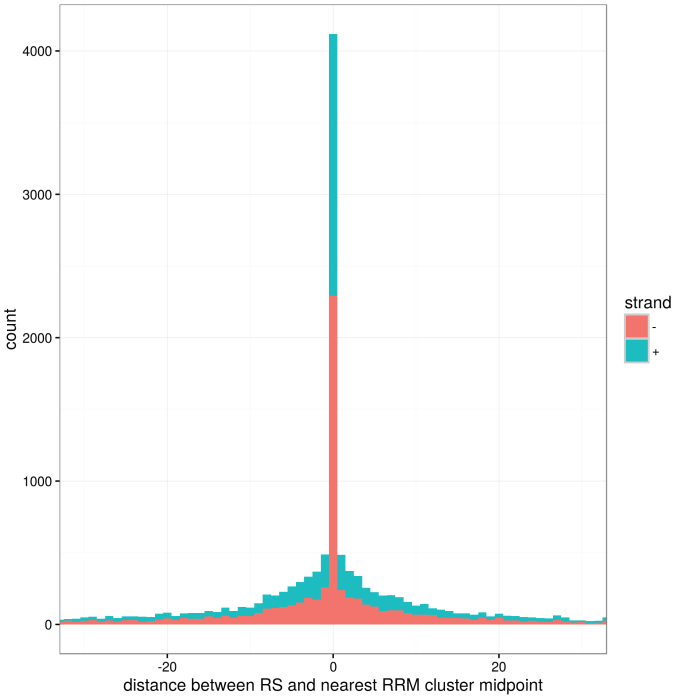


Figure S10
