## Supplemental Figure 6 for "The structure of yeast Npl3 bound to RNA reveals a cooperative sequence-specific recognition and an RNA chaperone role in splicing"

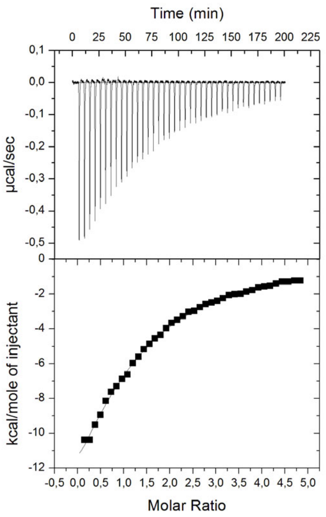


RRM1/2 R126A + AUCCAGUGGAA
Kd = 13 µM

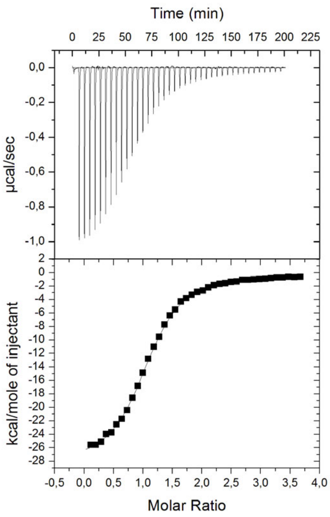


RRM1/2 R130A + AUCCAGUGGAA
Kd = 1.1 µM

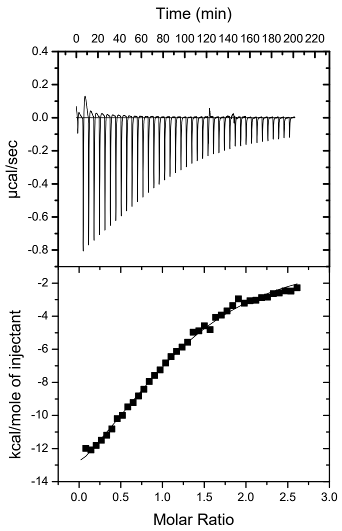


RRM1/2 F128A + AUCCAGUGGAA
Kd = 12 µM

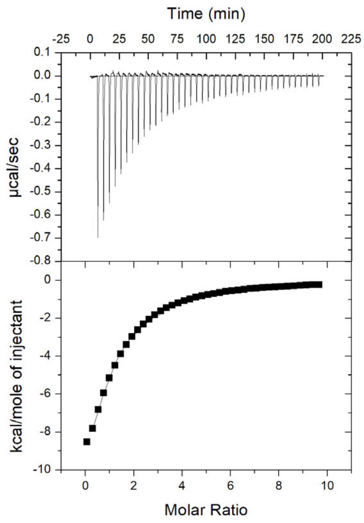


RRM1/2 F128AF160A + AUCCAGUGGAA
Kd = 12 µM

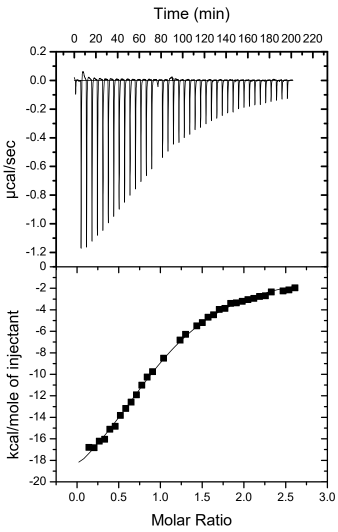


RRM1/2 K217A + AUCCAGUGGAA
Kd = 7 µM

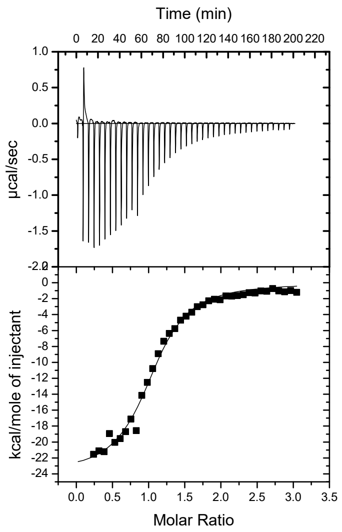


RRM1/2 Q214A + AUCCAGUGGAA
Kd =1.5 µM

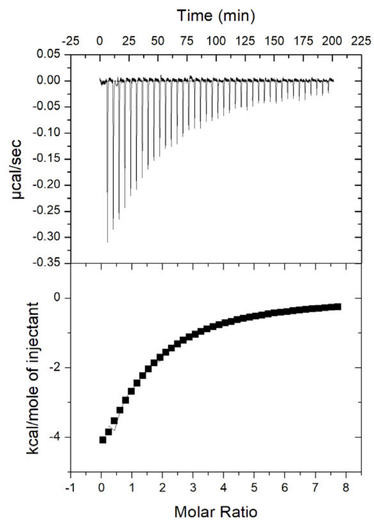


RRM1/2 Q214AK217A + AUCCAGUGGAA
Kd =16 µM

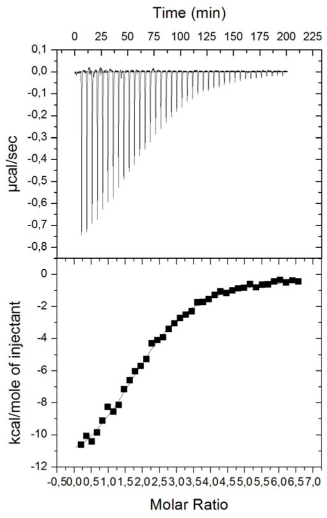


RRM1/2 F229A + AUCCAGUGGAA
Kd = 3.8 µM

Figure S6
