## Supplemental Figure 7 for "The structure of yeast Npl3 bound to RNA reveals a cooperative sequence-specific recognition and an RNA chaperone role in splicing"

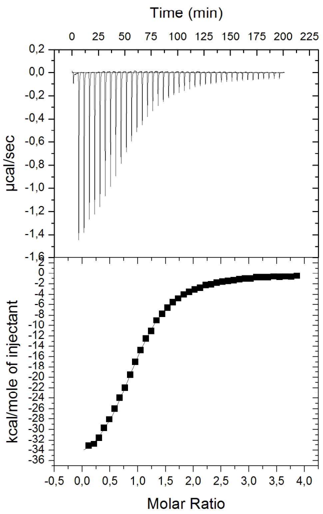


RRM1/2 + AUUCAGUGGAA
Kd = 1.3 μM

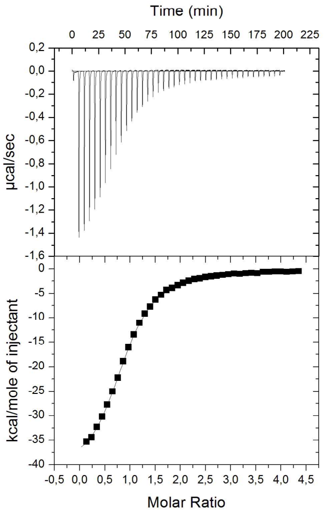


RRM1/2 + AUCUAGUGGAA
Kd = 1.2 μM

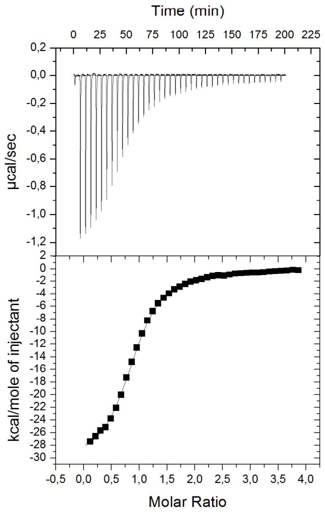


RRM1/2 + AUCCUGUGGAA
Kd = 0.87 μM

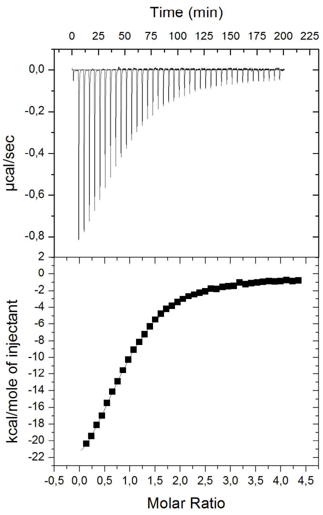


RRM1/2 + AUCCAAUGGAA
Kd = 2.7 μM

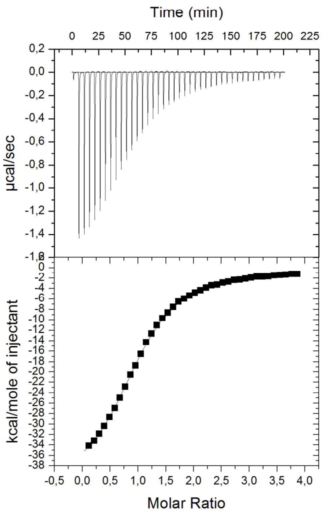


RRM1/2 + AUCCAGUAGAA
Kd = 2 μM

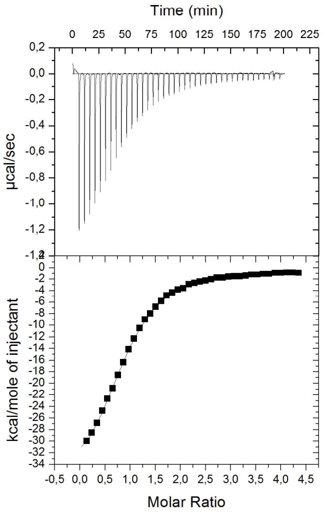


RRM1/2 + AUCCAGUGAAA
Kd = 1.95 μM

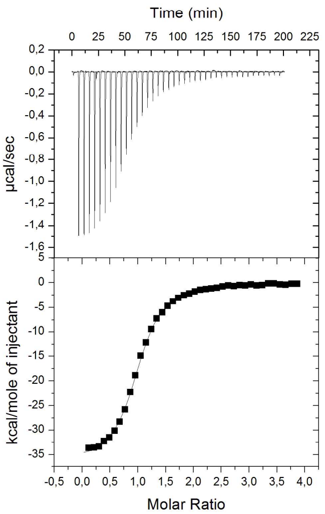


RRM1/2 + AUCC-GUGGAA
Kd = 0.5 μM

Figure S7
