## Supplemental Figure 11 for "The structure of yeast Npl3 bound to RNA reveals a cooperative sequence-specific recognition and an RNA chaperone role in splicing"

**A**

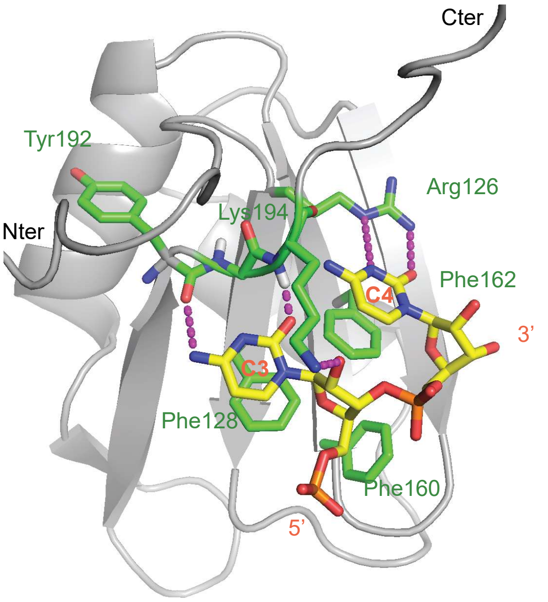


Npl3 RRM1 +
5′-AUCCAA-3′ RNA

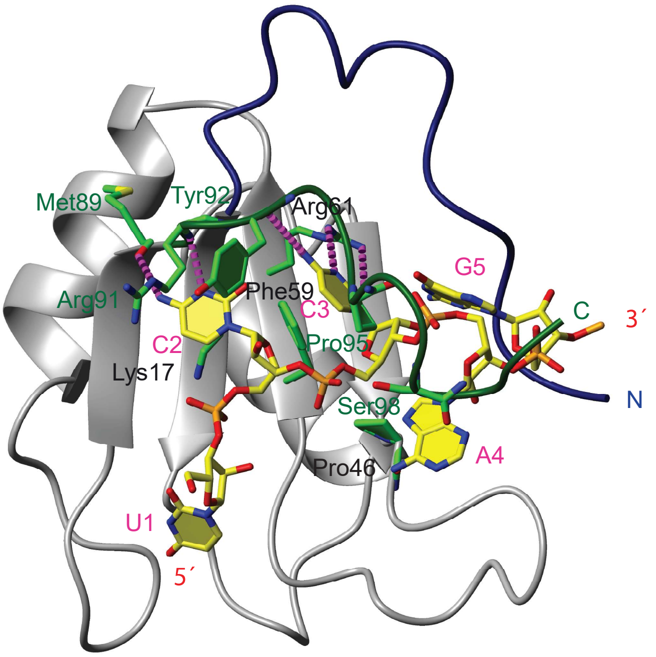


SRSF2 RRM +
5′-UCCAGU-3′ RNA

**B**

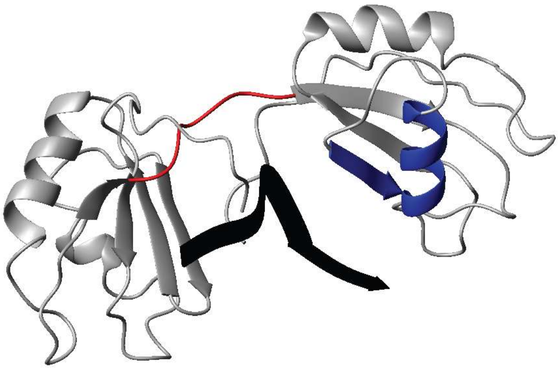

Npl3 RRM1/2
bound to RNA

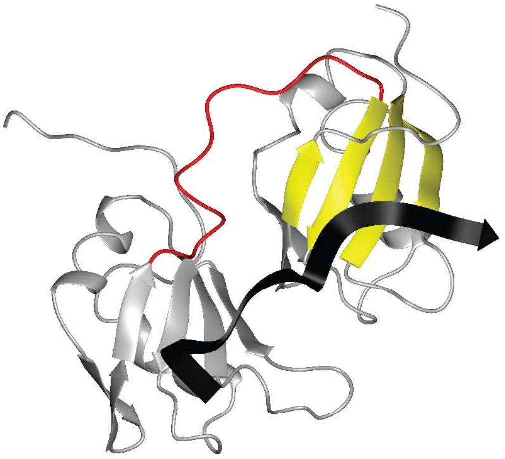

TDP43 RRM1/2
bound to RNA

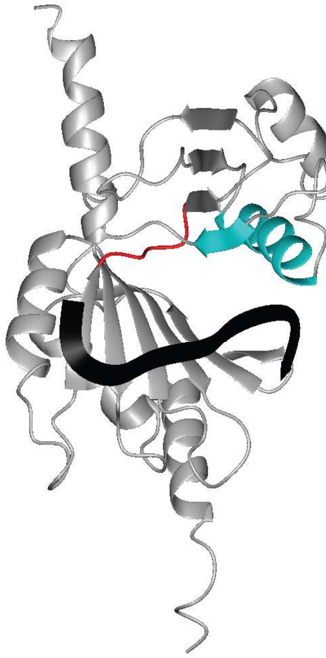

Dnd1 RRM1/2
bound to RNA

Figure S11
