## Supplemental Figure 12 for "The structure of yeast Npl3 bound to RNA reveals a cooperative sequence-specific recognition and an RNA chaperone role in splicing"

**A**

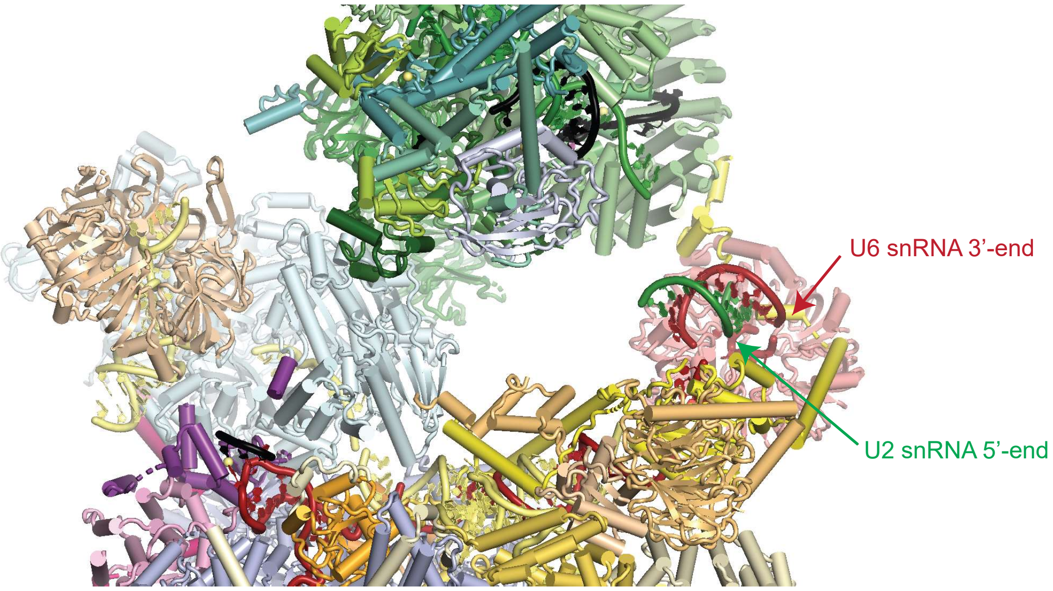


Spliceosome Yeast B complex

**B**

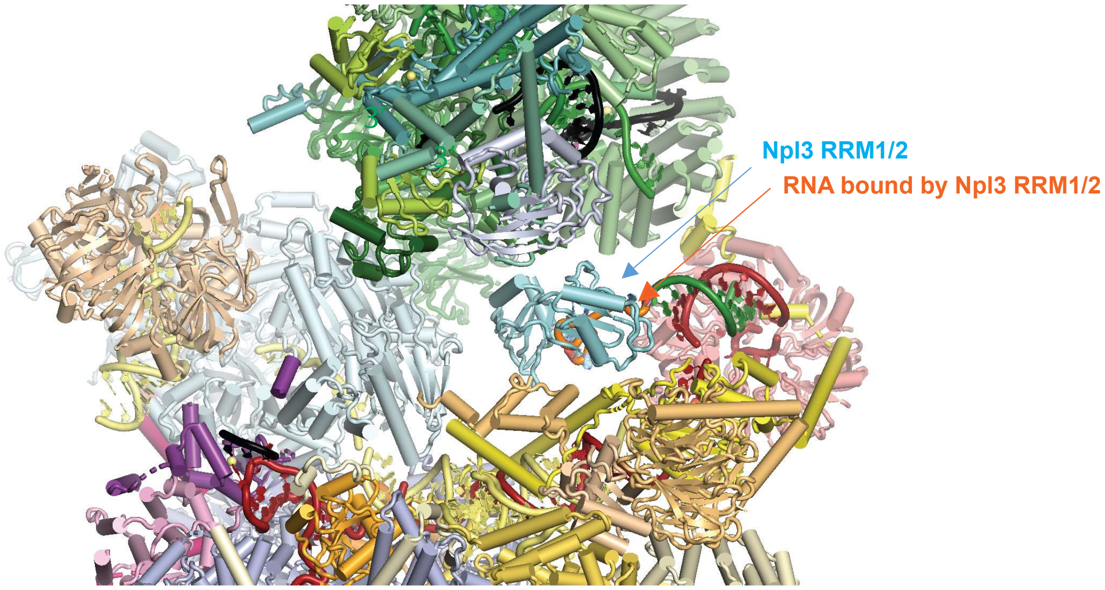


Spliceosome Yeast B complex after positioning Npl3 RRMs bound to RNA on the cryo-EM structure

Figure S12
