## Supplemental Table1 for "The structure of yeast Npl3 bound to RNA reveals a cooperative sequence-specific recognition and an RNA chaperone role in splicing"

### Table S1: Structural statistics of the Npl3 RRM1 in complex with AUCCAA RNA

| | Protein | RNA |
|---|---|---|
| **NMR distance and dihedral constraints** | | |
| Distance restraints | | |
| Total NOE | 2475 | 58 |
| Intra-residue | 463 | 40 |
| Inter-residue | | 18 |
| Sequential ($\|i-j\| = 1$) | 629 | 0 |
| Nonsequential ($\|i-j\| > 1$ ) | 1359 | 0 |
| Hydrogen bonds | 24 | 0 |
| Protein–RNA intermolecular | 135 | |
| Total dihedral angle restraints | | 6 |
| Protein | | |
| $\phi$ | 0 | |
| $\psi$ | 0 | |
| Nucleic acid | | |
| Base pair | | 0 |
| Sugar pucker | | 6 |
| Backbone | | 0 |
| Based on A-form geometry | | 0 |
| **Structure statistics** | | |
| Violations (mean and s.d.) | | |
| Number of distance constraints (>0.3 Å) (Å) | 6.3 ± 2.2 | |
| Dihedral angle constraints (º) | 0 | |
| Max. dihedral angle violation (º) | 0 | |
| Max. distance constraint violation (Å) | 0.43 | |
| Deviations from idealized geometry | | |
| Bond lengths (Å) | 0.0042 ± 0.0001 | |
| Bond angles (º) | 1.228 ± 0.016 | |
| Average pairwise r.m.s. deviation** (Å) | | |
| Protein | | |
| Heavy | 0.41 ± 0.06 | |
| Backbone | 0.14 ± 0.02 | |
| RNA | | |
| All RNA heavy | | 0.30 ± 0.05 |
| Complex | | |
| Protein and RNA heavy | | 0.41 ± 0.06 |

** Protein r.m.s. deviation was calculated using residues 125 to 195 for the ensemble of 10 refined structures. RNA r.m.s. deviation was calculated using nucleotides 110 and 112 for the ensemble of 10 refined structures.
