## Supplemental Table2 for "The structure of yeast Npl3 bound to RNA reveals a cooperative sequence-specific recognition and an RNA chaperone role in splicing"

**Table S2: Structural statistics of the Npl3 RRM12 in complex with AUCCAGUGGAA RNA**

| | Protein | | RNA |
|---|---|---|---|
| **NMR distance and dihedral constraints** | | | |
| Distance restraints | | | |
| Total NOE | 3788 | | 63 |
| Intra-residue | 728 | | 47 |
| Inter-residue | | | |
| Sequential ($\|i-j\| = 1$) | 1014 | | 16 |
| Nonsequential ($\|i-j\| > 1$ ) | 2008 | | 0 |
| Hydrogen bonds | 38 | | 0 |
| Protein–RNA intermolecular | | 189 | |
| Total dihedral angle restraints | | | 11 |
| Protein | | | |
| $\phi$ | 0 | | |
| $\psi$ | 0 | | |
| Nucleic acid | | | |
| Base pair | | | 0 |
| Sugar pucker | | | 11 |
| Backbone | | | 0 |
| Based on A-form geometry | | | 0 |
| **Structure statistics** | | | |
| Violations (mean and s.d.) | | | |
| Number of distance constraints (>0.3 Å) (Å) | 5.4 | | |
| Dihedral angle constraints (º) | 0 | | |
| Max. dihedral angle violation (º) | 0 | | |
| Max. distance constraint violation (Å) | 0.58 | | |
| Deviations from idealized geometry | | | |
| Bond lengths (Å) | | | |
| Bond angles (º) | | | |
| Average pairwise r.m.s. deviation** (Å) | | | |
| Protein | | | |
| Heavy | 1.23 ± 0.24 | | |
| Backbone | 0.96 ± 0.27 | | |
| RNA | | | |
| All RNA heavy | | | 1.42 ± 0.41 |
| Complex | | | |
| Protein and RNA heavy | | | 1.15 ± 0.22 |

** Protein r.m.s. deviation was calculated using residues 126 to 278 for the ensemble of 10 refined structures. RNA r.m.s. deviation was calculated using nucleotides 105 to 112 for the ensemble of 10 refined structures.
